# Telomere Position Effect Over Long Distance acts as a genome-wide epigenetic regulator through a common *cis*- element

**DOI:** 10.1101/2022.09.30.510336

**Authors:** Raphaël Chevalier, Victor Murcia Pienkwoski, Nicolas Jullien, Leslie Caron, Frédérique Magdinier, Jérôme D. Robin

## Abstract

Among epigenetic modifiers, telomeres, represent attractive modulators of the genome in part through position effects. Telomere Position Effect – Over Long Distances (TPE-OLD) modulates genes expression by changes in telomere-dependent long-distance loops, with a reach of 10Mb from a telomere. However, TPE-OLD remains poorly defined. To gain further insights into the genome-wide impact of telomere length on genomic and epigenomic regulation through TPE-OLD, we used cells with controlled telomere length combined to a genome wide transcriptome and methylome analysis. By integrating omics data, we identified a common *cis*-acting motif that behaves as an insulator or enhancer. Using reporter assays integrating this element, we uncovered the *trans* partners regulating this activity. Further exploiting our cellular model, we observed the depletion of one candidate factor, RBPJ, at TPE-OLD associated loci upon telomere shortening. We concluded that, at the genome-wide level, TPE-OLD is relayed by RBPJ binding Alu-like elements to telomeres that acts as enhancers. In response to external stimuli (*i*.*e*., Aging), TPE-OLD might act by coordinating telomere length to the action of Alu newly evolved enhancers in association with RBPJ.

## Introduction

Telomeres are unique nucleoprotein complexes composed by a repetitive 5’-(T_2_AG_3_)_n_-3’ motif associated to a protein complex named shelterin^1^. In vertebrates, a total of six proteins (*e*.*g*., TRF1, TRF2, POT1, RAP1, TPP1 and TIN2) characterize this structure located at the extremity of all chromosomes. Telomeres are also associated with other components: a long non-coding RNA known as TERRA (TElomeric Repeat-containing RNA)^2,3^ and a trimeric nucleoprotein complex (CST, composed of CTC1, STN1 and TEN1) binding to single-stranded telomeric DNA (telomeric overhang) where it is involved in chromosome end capping and telomere length regulation^4^. The-end assembled telomeric structure ensures genomic stability and prevents unwanted activation of the DNA damage response (DDR) signaling and repair of blunt ends^5–7^. At each round of replication, the DNA component of telomeres shortens, in all normal tissues regardless of their proliferative rate^8,9^, to a stage (*i*.*e*., short telomere length) preventing the protection mediated by shelterin units. Without this safeguard protective mechanism, cells enter replicative arrest, senescence or apoptosis^10,11^.

However, telomeric proteins are not solely restricted to the afore mentioned elements. Because telomere dysregulation is one of the main hallmarks of aging^12^ and due to its implication in cancer (re-activation of telomerase, alternative lengthening)^13,14^, recent works attempted to explore the full identity of telomere-associated proteins^15–17^. The telomere proteome is gradually expanding with more than 200 proteins^15,17^ involved in their functions including novel telomeric double strand DNA (dsDNA) binding proteins such as RBPJ^18^, HOT1^19,20^ and TZAP^21,22^. Recombination Signal Binding Protein For Immunoglobulin Kappa J Region, known as RBPJ or CSL, plays a central role in Notch signaling (*e*.*g*., pathways of cell fate-decision)^23,24^ and was recently involved in telomere homeostasis by directly binding to the telomeric dsDNA while recruiting factors preventing chromosome end fusion^18^. The homeobox telomere-binding protein 1, HOT1, was identified in a proteomic high throughput screen targeting telomerase recruiters^19^. HOT1 acts as bridge between telomere and active telomerase, tethering the telomere-elongating enzyme to its target and allowing its processing. Last, TZAP for telomeric zinc finger-associated protein, has been recently described as a telomere trimming protein, associated to long telomeres (with low density of shelterin units), thus setting the upper limit of telomere length^21,22^.

Altogether, current studies revealed that regulation of telomere homeostasis is dynamic and complex. Besides, the direct consequences of telomere shortening prior to cell cycle arrest, which represent but a short event in term of telomere erosion^25^ (when compared to the telomere shortening through life), remain to be characterized.

Among telomere-dependent processes, Telomere Position Effect (TPE), first described in *Saccharomyces cerevisiae*, is defined as the spreading of epigenetic marks spanning from the telomeres to adjacent genes leading to their silencing^26^. TPE triggers a proportional continuous silencing of genes depending on the distance from telomeres and telomere length^27^. Since then, this position effect has been described in human with only one gene detected so far *in vivo, DUX4*^28^. More recent findings, reminiscent of observations made in yeast, report the existence of another type of position effect dubbed TPE-Over Long Distances (TPE-OLD)^29^. TPE-OLD involves the formation of telomere length-dependent chromatin loops encompassing telomeres and subtelomeric genes. If TPE is restricted to a distance of a few kilobases (kb) from telomeres, the princeps description of TPE-OLD identified genes within 10 Megabases (Mb) from telomeres whose expression is directly impacted by telomere length, without any restrictions towards modulated expression (either up or down) by opposition to the repressive mechanism of TPE. Despite its identification and recent studies suggesting the genome-wide influence of telomere^29–32^, TPE-OLD features remain only partly characterized.

Here we sought to investigate the molecular basis of TPE-OLD via high throughput transcriptomic and epigenetics analyses using the same model that lead to the discovery of TPE-OLD^29^. We gained a more complete understanding of the genome-wide impact of TPE-OLD by identifying a common DNA motif shared by genes and DNA methylated region modulated by telomere length across cell types. This *cis* element act as an insulator/enhancer and shares association with telomere-associated proteins. Our results broaden our view of TPE-OLD at the genomic scale, opening the path to unthought-of implication of telomeres in aging and pathologies.

## Results

### Changes in methylation but not transcription is enriched at subtelomeres upon telomere shortening

To uncover the molecular basis of TPE-OLD, we applied the strategy reported in earlier studies^28,29,33^. Briefly, cells are immortalized with a floxable *hTERT* cassette and an additional CDK4 construct when required (*i*.*e*., myoblasts). Immortalized cells are then grown further to homogenize telomere length across chromosome ends (92 ends) with *hTERT* removed at various time points (reported by black triangle, Fig.1A) to generate isogenic clones with controlled telomere length. Importantly, all cells roughly spend the same time in culture, thus minimizing effects imputable to cell culture conditions^34,35^. We applied this strategy to fibroblasts, myoblasts and their differentiated counterpart, myotubes.

**Figure. 1.**
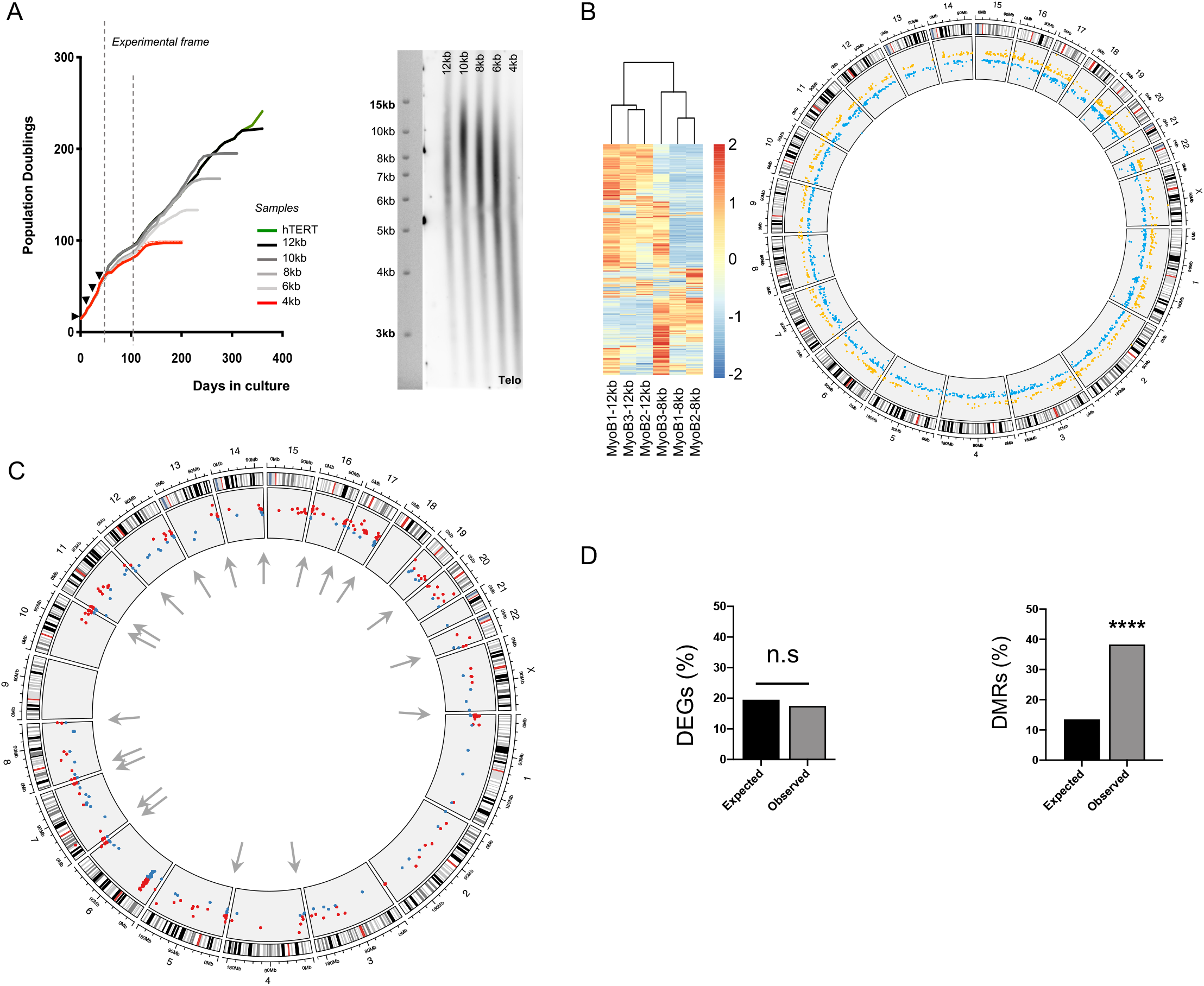
DNA methylation changes are enriched at subtelomeres upon telomere shortening. **A**. Growth curve and associated Telomere Restriction Fragment analysis (TRF) of the cellular model used for the study. Time point of hTERT excision is symbolized by black triangle along with the frame used for experiments by dashed lines. Isogenic clones, reported with their estimated telomere length were explored before any sign of DNA damage (supplemental figure 1). **B**. Heatmap of unsupervised clustering of differentially expressed genes (DEG) in isogenic myoblasts clones with long (12kb) and shorter telomere (8kb) along with a Circos’ representation of DEGs distribution across the genome. **C**. Repartition of differentially methylated regions (DMRs) across the genome in isogenic myoblast clones with long (12kb) and shorter telomere (8kb). Arrows (grey) point to DMRs located at subtelomeres. **D**. Proportion of expected DEGs in subtelomeric regions compared to observed DEGs (left panel) along with the proportion of expected DMRs in subtelomeric region compared to observed DMRs (right panel) in our dataset. Only subtelomeric DMRs are statistically enriched. Geometric test (DEGs), *X*_2_ test (DMRs); α = 0.05. *p* **** <0,0001.

Next, we estimated the mean telomere length and exploited cells with long and shorter telomeres for each cell type (Fig. 1A, Supplemental 1A). For myoblasts and myotubes, we used isogenic clones with an average telomere length of 12; 10; 8; 6 or 4kb and 14 or 8kb for fibroblasts. Of note, DNA damage signals detected with γH_2_AX staining were significant only in myoblasts with the shortest average telomere length (4kb, Supplemental 1B-C). Then, we performed a transcriptomic analysis in myoblasts with long (12kb) and shorter telomeres (8kb) along with a methylome analysis using an EpicArray approach in myoblasts with long and shorter telomeres (12; 10; 8; 6; 4kb; respectively). Cells with an average telomere length of 4kb were included as a cell ‘crisis’ time point and were removed from later observations, unless explicitly mentioned.

From the transcriptomic analysis, differentially expressed genes (DEGs) were found across all chromosomes and allowed the clustering of myoblasts in agreement with their average telomere length (Fig. 1B). Regarding biological pathways enriched in myoblasts with shorter telomeres, we note that terms associated with upregulated genes are related with muscle fusion, function and contraction whereas terms associated with downregulated genes are involved in cell migration (Supplemental Fig. 2). These results suggest that shorter telomere length might act as an effector triggering premature muscle differentiation in myoblasts.

Methylation analysis of 850K CpGs across the genome revealed an overall trend towards hypermethylation in cells with shorter telomeres (Supplemental Fig. 3), without overrepresentation of specific regions (*e*.*g*., CpG island) nor functional elements (*e*.*g*., 3’UTRs, TSS). A trend consistent with previous reports^36^. Similar to transcription results, we report several Differentially Methylated Regions (DMRs) across the genome (Fig. 1C). By filtering out DMRs that were not consistent across myoblasts with decreasing telomere length (*e*.*g*., constant hyper/hypomethylation, constant DMR position), we observed a fair proportion of DMRs located in subtelomeric regions (grey arrow, Fig. 1C).

We confirmed our observation by reporting the expected percentage of DEGs (corresponding to the proportion of genes located in subtelomeres; *i*.*e*., 20%) or DMRs compared to the result obtained from our transcriptomic and methylome analysis (Fig. 1D, geometric test, *p*=0.99; *X*_2_ test, *p<*0.0001; respectively). Besides, we report the same results regarding DNA methylation in two additional cell types (*i*.*e*., myotubes, fibroblasts; Supplemental Fig. 4). Hence, DMRs associated to telomere length are enriched in subtelomeres, suggesting that telomere shortening first impacts DNA methylation.

### TPE-OLD acts as a genome-wide epigenetic regulator through a unique DNA motif

Next, we sought for a common signature by crossing the transcriptome and methylome analysis using comparison between DEGs and DMRs. We found a 35 bp DNA sequences shared by both datasets (Fig. 2A). We dubbed this motif “WE” for Wide Effect motif.

**Figure. 2.**
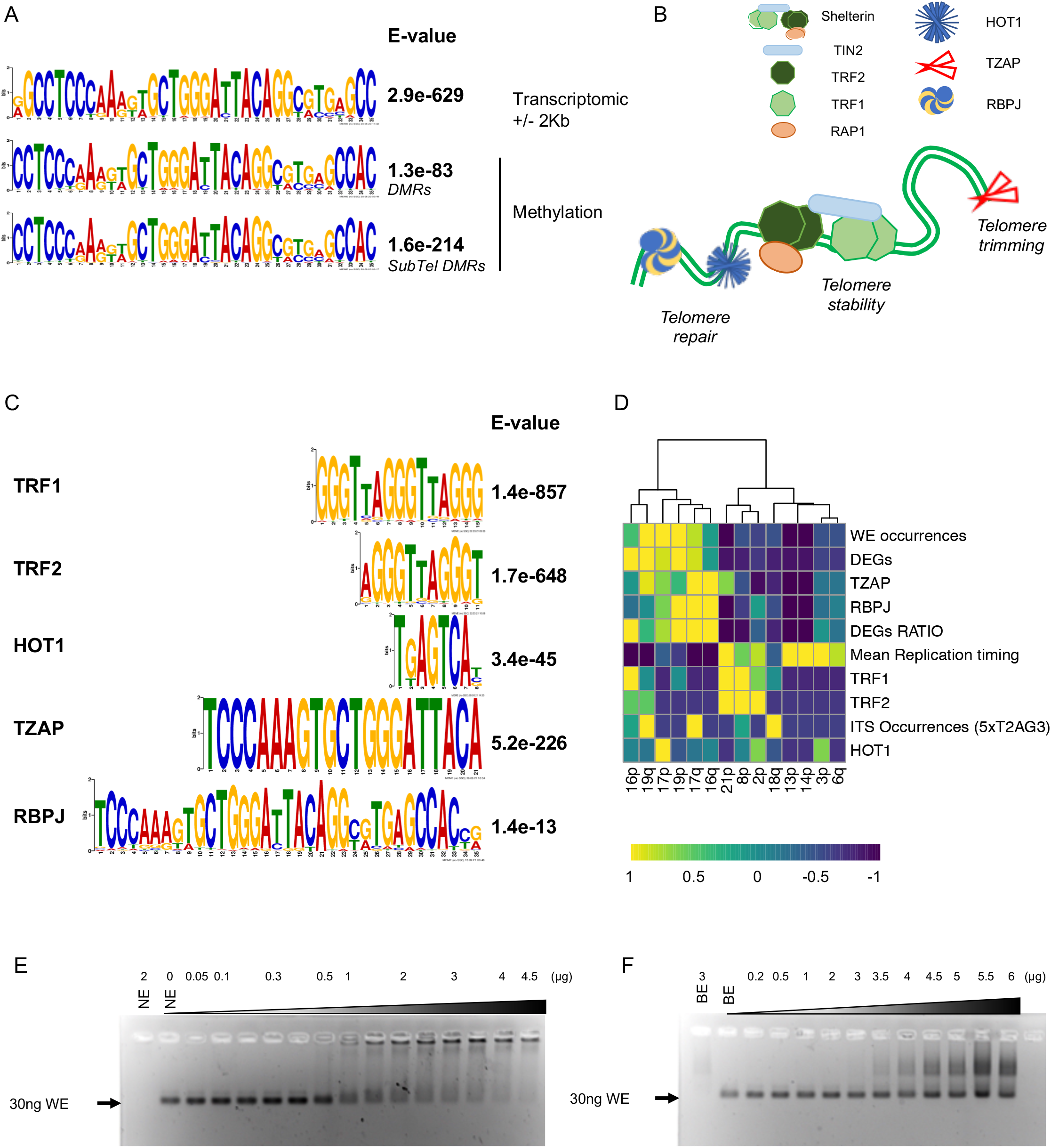
Transcriptomic and DNA methylation changes upon telomere shortening are associated to a unique DNA motif. **A**. Motif logos of the top sequence present in direct proximity of either DEGs (± 2kb), DMRs or DMRs located in subtelomeres from data collected in isogenic clones with long (12kb) and shorter (8kb) telomeres. For each motif, we report the associated E-value. **B**. Graphical representation of the main telomere-associated proteins binding to dsDNA and their main involvement in telomere homeostasis. **C**. Motif logos and associated E-values of the top sequence present in previously published ChIP-Seq experiments. We report the most abundant motif associated with each dsDNA binding telomeric associated proteins. **D**. Unsupervised hierarchical clustering (WardD2; Manhattan distance) of specific subtelomeres (10Mb) with motif occurrences (WE), DEGs and telomeres associated factors (replication timing, proteins, Internal Telomeric Sequences ITS). Parameters were extracted from available data set and normalized to a mean of 1. We report DEGs as total DEGs localized at each subtelomeres and DEGs ratio when corrected for the gene density (Complete set is reported in Supplemental figure. 5). **E-F**. Electromobility Shift Assay performed in a 2% agarose gel. For each assay 30ng of the 35bp motif was loaded and run with increasing amount of either nuclear (**E**) or bacterial (**F**) extract. Shift is only visible in the experiment using nuclear extracts.

Stemming from previous work describing the influence of telomeres over the genome^29,32,37,38^ and taking advantage of available datasets (ChIPSeq)^19,39,40^, we compared the WE sequence to the main motifs enriched in dsDNA binding proteins associated to telomeres; each with a described role in telomere homeostasis^19^ (Fig. 2B). As predicted the main motif associated to the shelterin proteins (TRF1, TRF2) correspond to the canonic telomere sequences T_2_AG_3_ and HOT1 to a small 5’-TGAGTCA-3’ motif. Singularly, TZAP and RBPJ two proteins associated with telomere trimming and repair presented an enrichment for a motif highly similar to WE (Fig. 2C).

To further investigate the WE sequence with regard to telomeres homeostasis, we explored potential associations between various factors including proteins; WE occurrences; DEGs and DEGs ratio (normalized to the total number of genes present at telomere); Internal Telomeric Sequences (ITS) and replication timing^41,42^, respective to chromosome ends (Supplemental Fig. 5A). Unsupervised clustering, at the chromosome-end level, confirmed the correlation between WE, RBPJ and TZAP along with an inversed relationship with replication timing. Additionally, because telomeres are restricted to eukaryotes, we tested the specificity of WE by EMSA using either nuclear (Fig. 2E) or bacterial (Fig. 2F) extracts. We observed that DNA-protein complexes were specific and restricted to assays using nuclear extracts (Fig. 2E, Supplemental Fig. 5B).

Altogether our data suggest a common *cis* motif present in TPE-OLD genes associated with TZAP and RBPJ enrichments and inversely proportional to the speed of telomere replication with earlier replicative telomeres holding the most the most WE occurrence.

### TPE-OLD motif signature displays an enhancer/ insulator activity

To gain insight on the direct properties linked to WE in gene regulation (*i*.*e*., enhancer, repressor, insulator), we took advantage of reporter constructs inserted in a controlled chromatin context^43,44^. Briefly, our system includes a Hygromycin resistance gene and consists on an eGFP cassette driven by a pCMV promoter followed by a motif insertion site. Thus, integration into the genome (mostly in a heterochromatin context) allows one to test the proprieties associated to the inserted sequences. We tested different WE-derived motifs each: WE, Scramble, reversed orientation and mutations (Fig. 3A). Further, we used an analog construct, pCMVTelo, carrying a telomere seed forcing the insertion in a telomeric position after a telomere crisis and healing at any chromosome ends^43^. After control of single insertion in HEK 293 cells by DNA FISH (Fig. 3B), we maintained all cell populations under Hygromycin selection and recorded their eGFP expression profiles (intensity, cell proportions; Fig. 3C, Supplemental Fig. 6). We report an increase of eGFP positive cells restricted to cells carrying the WE sequence (Mean± SEM= 61.31± 4.35) in comparison of all others (*p*<0.0001, Holm-Sidak’s multiple comparisons test). As expected, counterpart constructs in a telomeric context (pCMV, pCMVTelo; respectively) exhibited a decrease of eGFP-positive cells but for the EW motif. Our results suggest that WE hold proprieties similar to insulators and enhancers. In a reversed orientation, dubbed EW, this motif contains a T_2_AG_3_ sequence and seems to act as a silencer.

**Figure. 3.**
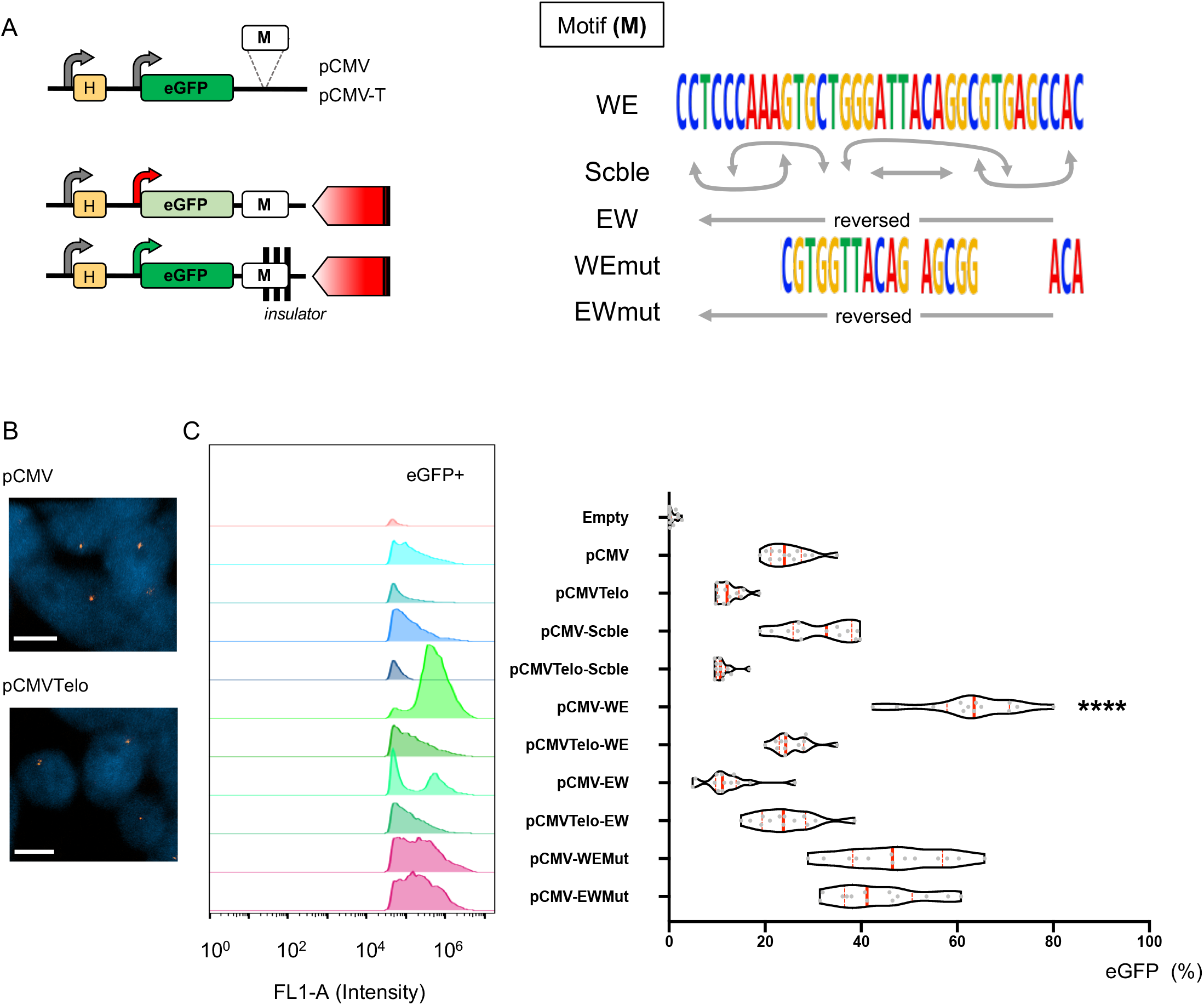
The Associated TPE-OLD motif presents an enhancer/insulator activity. **A**. Graphical representation of the construct (left) and DNA sequences used (right) for reporter assays. Briefly sequences to test are inserted at a multiple restriction site (M) located directly after the *eGFP* reporter gene and upstream of a telomere seed for pCMV-T. Each construct (WE; Scble; EW; WEMut; EWMut) is transfected and populations of cells selected in the presence of Hygromycine (H). Constructs allow one to test either Position Effects (pCMV) or Telomere Position Effect (pCMV-T) and the insulator-related proprieties of the inserted motif. Condition are optimized for a single insertion per cell. **B**. Representative images showing single insertion of constructs (pCMV; pCMV-T) in HEK 293T cells. **C**. eGFP expression profile and corresponding quantifications determined by flow cytometry in cells transfected with various constructs. After transfection of constructs and selection, eGFP levels were measured by flow cytometry for 3 consecutive weeks. Assays were performed in quadruplicate for a total of n=12. We report all data points in box and violins. Medians and quartiles are shown (red and dashed lines, respectively). Holm-Sidak’s multiple comparison test; α = 0.05. *p* **** <0,0001.

### WE activity can be modulated by RBPJ and SMCHD1

To challenge our hypothesis regarding WE associated proprieties, and its role in the TPE-OLD mechanism, we tested a subset of WE-gene motifs. Thus, we picked WE-related sequences found in the vicinity of DEGs from our transcriptomic analysis (Fig. 1) such as *C16Orf74* (Motif length: 201bp; distance from gene promoter: 22kb); *BET1L* (195bp; 1.2kb); *GIPC3* (372bp; 2.5kb); *GLIS2* (413bp; 55kb) and inserted these gene-related motifs in both constructs (pCMV, pCMVTelo).

All motifs were associated with an increased in eGFP-positive cells (Fig. 4A, Supplemental Fig. 7). Some WE-gene motifs (*BET1L, GLIS2)* behaved equally as WE with a decrease signal at telomeric positions (pCMVTelo) whereas two others (*C16Orf74, GIPC3*) did not exhibit significant changes depending on their chromatin insertion context. Our observations further advocate for an insulator/ enhancer role of WE with differences imputable to the variation respective to the WE motif found at different TPE-OLD genes, suggesting divergences in their overall regulations.

Next, we took advantage of our set system (*i*.*e*., pCMV/Telo in HEK 293) to evaluate possible *trans* partners associated with TPE-OLD using an siRNA approach. In agreement with earlier findings (Fig. 2D), we targeted TRF2, RBPJ and TZAP. Furthermore, we added CTCF due to its global role in chromatin organization and SMCHD1, an important chromatin remodeler found at telomeres in separate studies^15,16,45^. Validation of our siRNAs showed specific downregulations across condition (Supplemental Fig. 8). However, siRNAs targeting *TERF2* also significantly decreased *RBPJ*. As expected, no changes were observed in our control condition (pCMVScble; Supplemental Fig. 9). Strikingly, independent of the WE-related motif tested (WE, *GLIS2*), we observed a significant decrease of eGFP-positive cells upon either *TERF2* or *RBPJ* downregulation (Fig. 4B-D; Supplemental Fig. 10–12). Because si*TERF2* conditions also decreased *RBPJ*, one could hypothesize that effects observed in si*TERF2* are imputable to the off-target effect inducing *RBPJ* downregulation potentially caused by their cross-talk^18^. Concurring, changes were less pronounced in si*TERF2* than si*RBPJ* situations across constructs (Fig. 4B-D). To note, we report a lower proportion of eGFP positive cells upon *SMCHD1* downregulation, independent of constructs and motifs (WE, *GLIS2*). Taken together our observations confirmed the insulator/enhancer properties of WE (and associated sequences) and support RBPJ and SMCHD1 as putative *trans* effectors, whereas TPE-OLD mechanism appears independent of TZAP or CTCF and only weakly associated with TRF2 for the different sequences tested.

**Figure. 4.**
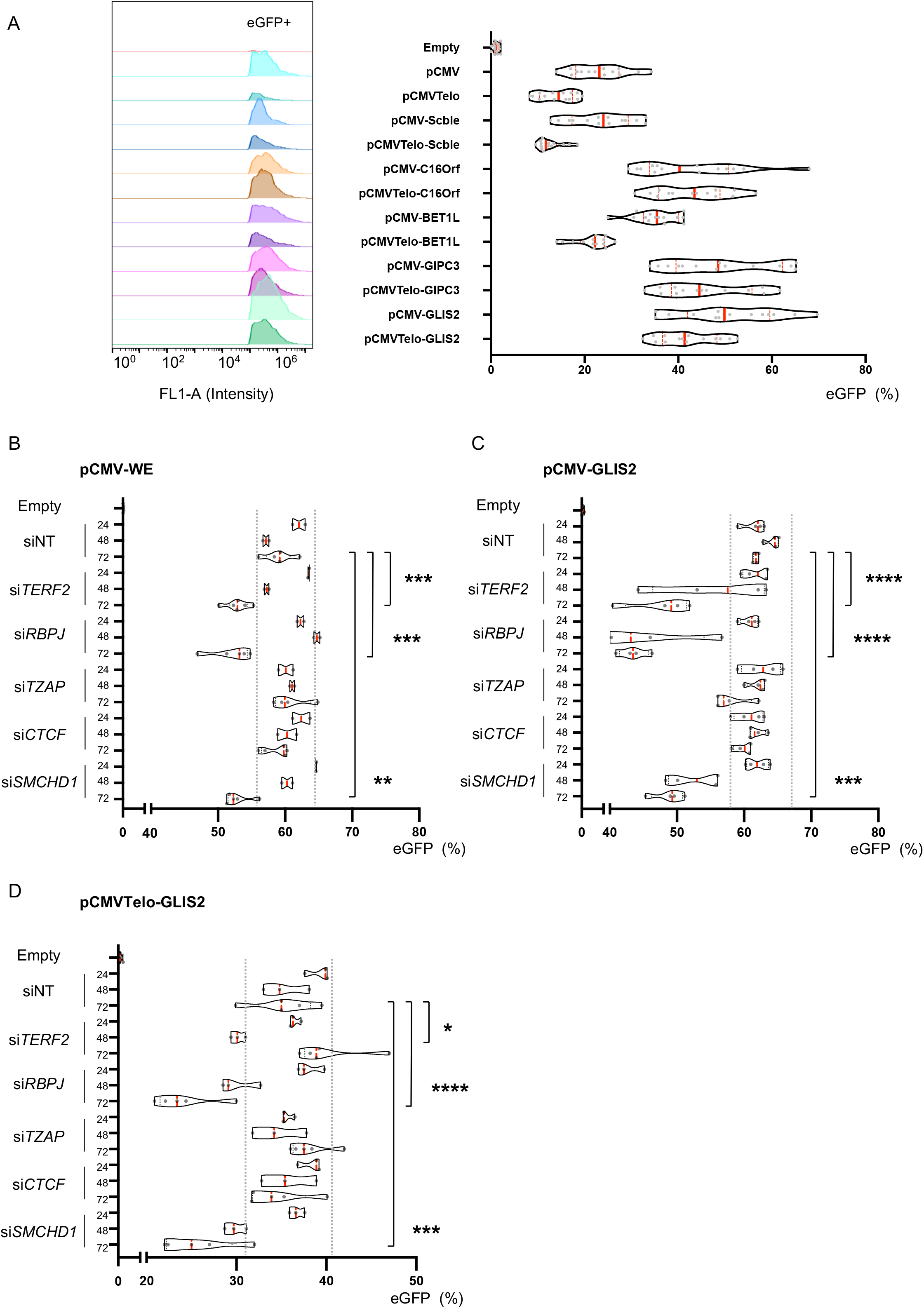
WE activity is modulated by RBPJ and SMCHD1 expression level. **A**. eGFP expression profile determined by flow cytometry in cells transfected with reporter genes containing the motif found in proximity of DEGs (*C16orf74, BET1L, GIPC3, GLIS2*) and corresponding quantifications. After transfection of constructs and selection, eGFP levels were measured by flow cytometry for 3 consecutive weeks. Assays were performed in quadruplicate for a total of n=12. We report all data points in box and violins. Medians and quartiles are shown (red and dashed lines, respectively). **B-D**. Quantification of reporter assay after transfection of siRNA targeting epigenetic modulators and dsDNA telomeric binding proteins. Cells with either pCMV-WE (B), pCMV-Glis2 (C) or pCMVTelo-Glis2 (D) constructs were transfected with siRNA and the proportion of eGFP positive cells was measured by flow cytometry at 24, 48 and 72 hours post transfection. For each condition, we report biological quadruplicate (n=4) in box and violins. Medians and quartiles are shown (red and dashed lines, respectively); dashed grey lines are shown to represent the variability associated with the NT condition. Holm-Sidak’s multiple comparison test; α = 0.05. *p* * <0,05; *p* ** <0,005; *p* *** <0,001; *p* **** <0,0001.

### TPE-OLD is regulated by RBPJ but not dependent on TRF2 or TZAP enrichment

To confirm our findings regarding the molecular signature of TPE-OLD, we further investigated its *cis* (WE) and *trans* (RBPJ, SMCHD1) partners in myoblasts with long and shorter telomeres (Fig. 1A). First, we noticed that WE and subsequent motifs found at proximity of TPE-OLD genes shared similarity with Alu repeats. Indeed, using our transcriptomic and methylome data (Fig. 1B-C), we observed that Alu repeats are enriched in proximity of DEGs (Supplemental Fig. 13); specifically, the sub class of Alu*Y*. Hence, we asked if DNA methylation of this particular repeat was modulated in our isogenic clones. Interestingly, we record a constant trend towards hypomethylation throughout our panel of cells with decreasing telomere lengths (*i*.*e*., 12 to 4kb). This trend is restricted to Alu as suggested by the unmodified DNA methylation profile of TAR1, the main known subtelomeric repeat^46^ (Fig. 5A, Supplemental Fig. 13).

**Figure. 5.**
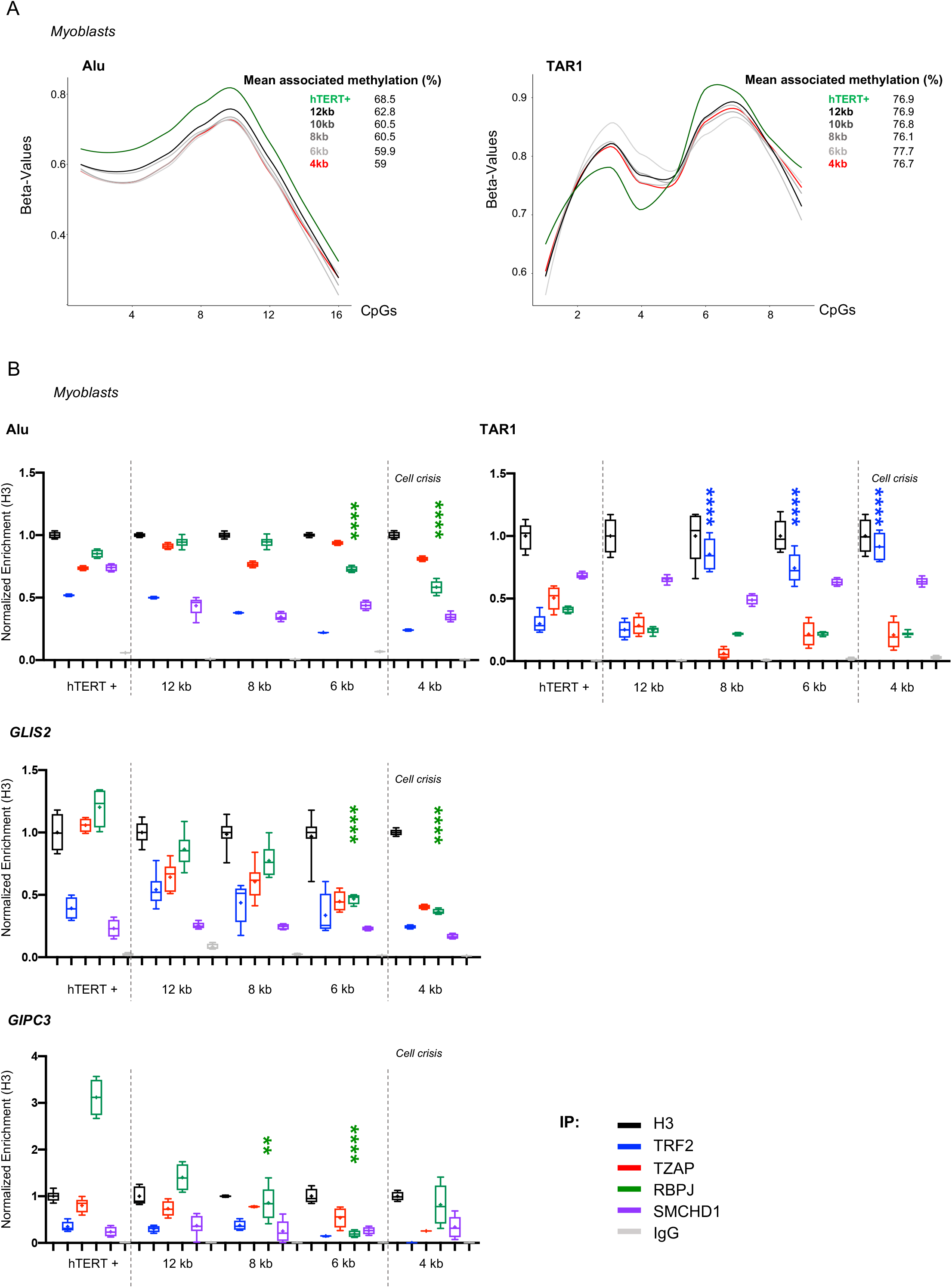
Associated TPE-OLD motif is regulated by RBPJ enrichment at specific loci *in vitro*. **A**. Smoothed mean methylation per CpG obtained after high throughput targeted bisulfite sequencing of either Alu or TAR1 repeats corresponding to amplicons of 219bp with 16 CpGs and 150bp with 9 CpGs, respectively. We report the methylation beta_value for all CpGs in the sequence of interest. A value of 1 corresponds to a fully methylated CpG; 0 to unmethylated. **B**. Protein enrichment at Alu, TAR1, *GLIS2* and *GIPC3* loci detected by ddPCR after ChIP in isogenic clones with various telomere length. We report the enrichment of H3 (black), TRF2 (blue), TZAP (red), RBPJ (green), SMCHD1 (purple) and IgG (grey) normalized to H3. Dashed grey lines are shown to represent the external limit of telomere length (active telomerase, 4kb; respectively). ChIP-ddPCR were performed in biological quadruplicate. Holm-Sidak’s multiple comparison test; α = 0.05. *p* * <0,05; *p* ** <0,005; *p* *** <0,001; *p* **** <0,0001.

Further, to confront our results from reporter assays in their true genetic and epigenetic context, we evaluated protein enrichments at different repeats (Alu, TAR1) and loci (*GLIS2, GIPC3, BET1L, C16Orf74*) in isogenic myoblasts with controlled telomere length, including cells with active telomerase and cells reaching crisis (positive to DNA damages; average telomere length: 4kb). We performed ChIP-ddPCR experiments after immunoprecipitation using either anti-H3, -TRF2, -TZAP, -RBPJ or -SMCHD1 antibodies; acknowledging the complete set of potential *trans* partners highlighted for TPE-OLD (Fig. 2,4). Regarding repeats, we found a significant decrease in both RBPJ and TRF2 enrichment at Alu-associated loci (Fig. 5B; *p* <0.0001 and *p*<0.0001; respectively, Holm-Sidak’s multiple comparisons test), reminiscent of changes reported for DNA methylation. In the context of TAR1-associated loci, we showed an enrichment of TRF2 in cells with shorter telomeres (8 to 4kb, *p* <0.0001; Holm-Sidak’s multiple comparisons test). This could be caused by the relocation of TRF2 at subtelomeres upon telomere shortening, as reported by others^32^.

At *GLIS2*, we noticed a constant trend towards depletion of RBPJ (*p* <0.0001; Holm-Sidak’s multiple comparisons test) whereas other proteins remained unchanged, with the exception of a significant depletion of TZAP (6kb, *p=* 0.044; Holm-Sidak’s multiple comparisons test). Moreover, we note a global enrichment of RBPJ and to a lesser extent, TZAP, in cells with active telomerase (labelled hTERT+). Similar observations were made at *GIPC3, BET1L* and *C16orf74* loci (Fig. 5B, Supplemental Fig. 14).

Our results described a dynamic enrichment at TPE-OLD loci of RBPJ proportional to telomere length whereas others (TZAP, SMCHD1) remained stable, with the exception of TRF2 that, in two separated contexts (TAR1, *BET1L*), is enriched in cells with short telomeres. This observation might be imputed to the close proximity of the loci (2-4kb, 200kb; respectively) to telomeres^32^. Taking together our results detailed the molecular basis of TPE-OLD through a common *cis* motif that can be directly impacted by RBPJ in *trans*.

## Discussion

In this study, we explored the mechanisms related to TPE-OLD, a progressive phenomenon closely linked to telomere shortening and thus aging^12^. Aging-related changes in gene expression include modifications of epigenetics marks^47,48^ and chromatin structure, both processes described in TPE-OLD but not further investigated^29,37,38^. Here, we found that TPE-OLD is defined by a *cis* element and proteins previously found at telomeres.

Our observations on the *cis* element showed a sequence similar to Alu elements and more precisely Alu*Y*, the most recent subclass of Alu^49^. The repartition of Alu in the genome is highly correlated with CG content, gene density and enriched in intragenic regions^50^, a seemingly odd observations if one considers Alu as inactive elements^51^. Consistent with our findings, Alu elements can form distant genomic interactions^52^ and modulate transcription by acting as a *cis* element for transcription factors^53^. However, if recent emerging studies report their enhancer activities^53^, Alu elements and their different subclasses are able to induce either up-or downregulation of associated genes^54^. This observation is also compatible with TPE-OLD^29^. In agreement with changes in DNA methylation in isogenic clones with long and short telomeres, Alu methylation level decreases with age^55^ and in pathological context such as Alzheimer^56^ (where telomere shortening has been observed), further supporting our hypothesis linking telomere length, TPE-OLD, DNA methylation and Alu elements. Enhancer-like characteristics of Alu, as the one evidenced by our work are shown to follow an evolutionary continuum^53^. Hence, “finalized” proto-enhancer corresponds to the most recently evolved Alu subclass, *i*.*e*. Alu*Y*; similar to our *cis* element found enriched at TPE-OLD associated genes and validated by a previous report^29^.

Beyond, our exploration of the WE *cis* element uncovered proteins associated to TPE-OLD. As shown by our siRNA approach, TPE-OLD *trans* partners can modulate its effect (*e*.*g*., TRF2, RBPJ and SMCHD1). However, their involvement is restrained to their associated genomic context. Indeed, TRF2 can act as a potential modifier of the *cis*-element *in vitro* but is not observed *in vivo* in the genomic context tested (*GLIS2*, Alu, TAR1) as we do not report any depletion of TRF2 at these loci. On the contrary, we showed an enrichment of TRF2 at TAR1, concomitant to telomere shortening. TRF2 is relocated towards the most telomeric sequences as telomere shortens, as previously suggested by others^32^. Likewise, regulation by SMCHD1 is possible but appears context-dependent, echoing previous work deciphering its role in Human^45,57,58^. Strikingly, the most consistent *trans* partner of TPE-OLD (*i*.*e*., acts on WE element and is depleted at TPE-OLD loci upon telomere shortening), RBPJ, is a transcriptional factor that was found to function by triggering recruitment of chromatin remodeling complexes, including histone modifiers^59^; suggesting a dynamic interaction with potential enhancers^60^. In our hands, a potential common enhancer interacting genome-wide with RBPJ might be WE. Of note, previous studies did not consider individual telomere length and only used overall average telomere length^25^ (*i*.*e*., masking outliers). This simplification does not integrate the well-accepted notion that human telomere ends (*e*.*g*., 92 chromosome ends) are highly variable from one chromosome to another and between respective arms for each individual^25,61^. Using our system, we minimized telomere variations in a homogenous genetic background (isogenic clones), hence allowing one to investigate TPE-OLD. One could hypothesize, that the combination of telomere length heterogeneity coupled with WE-polymorphisms (Alu*Y*-like repeats) offers an endless combination of systems where TPE-OLD associated genes are affected upon stimuli either by telomere shortening (progressive) or WE-polymorphisms (static). This new context could explain the enrichment of deregulation reported by others at subtelomeres^31^. In large panels, chances are that telomere shortening and WE-polymorphisms allowing genes to be freed from TPE-OLD are more consistently frequent at these specific loci.

Taken together, this indicates that telomeres and TPE-OLD impact has been largely overlooked in genome regulation as well as in diseases where telomere might act as a modifier such as age-related pathologies. This could also hold true for a number of diseases linked to subtelomeric imbalance where identification of TPE-OLD sensitive genes might help in solving the missing heritability or explain incomplete penetrance and phenotypical variability. Our descriptive work combining *in vitro* and *in silico* approaches opens new research perspectives of research by stressing the importance of TPE-OLD and its implication in human physio-pathology to evaluate diseases patho-mechanisms, disease risks and comorbidities associated to aging.

### Online Methods

Detailed methods and supplemental material can be found with this article online.

## Acknowledgments

We thank Dr. Nicolas Lenfant for helpful advices on bioinformatics analysis. The Genomic and Bioinformatic Core (GBiM) and Imaging Facility at MMG. The Dr. Christophe Picard and Dr. Pascal Pedini at EFS for access to ddPCR equipments. This study was funded by Aix Marseille University (A*Midex a French “Investissements d’Avenir programme” - pépinière d’excellence) and the INSERM-first steps program. This project has also received funding from the Excellence Initiative of Aix-Marseille University - A*Midex a French “Investissements d’Avenir programme”-Institute MarMaRa AMX-19-IET007 (VMP, fellowship). Study sponsors had no role in study design, collection analysis, data interpretation, writing of the report, or decision to submit the paper for publication.

## Declaration of interest

the authors declare no competing interest.

## Supplemental Figures

**Supplemental Figure 1.**
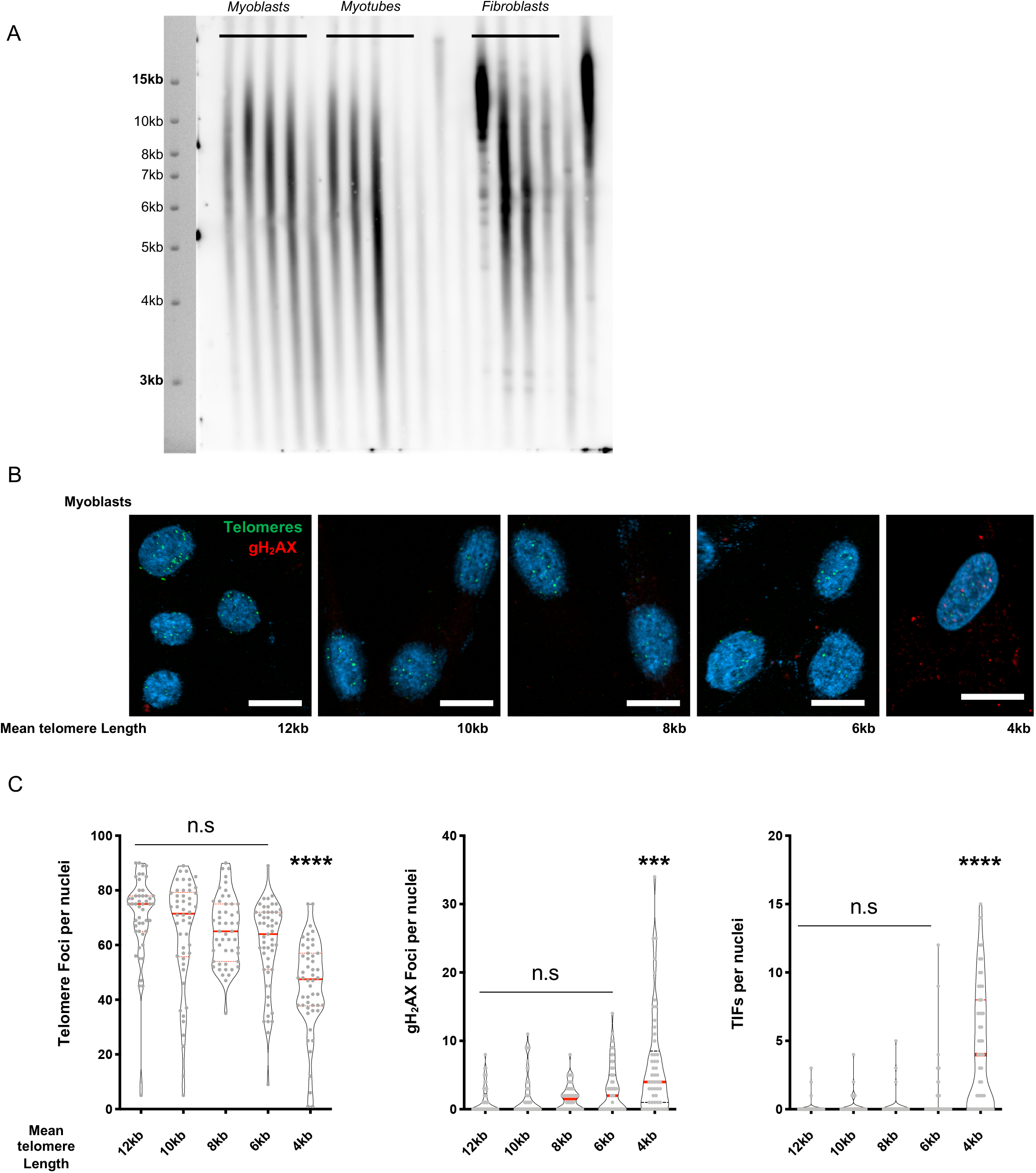
**A**. Representative Telomere Restriction Fragment analysis (TRF) of the cellular models used for the study as described in Stadler *et al*. 2013 and Robin *et al*. 2014. Myoblasts and their corresponding myotubes (post-differentiation) and fibroblasts were analyzed before detection of DNA damage signals **B**. Representative images of γH2AX and telomeric staining. Briefly Isogenic myoblast clones (described in Figure1.) with various telomere length were fixed, hybridized with a telomeric probe (C-Rich) and further stained using a γH2AX antibody. A scale of 10 µm is reported by a white bar in each frame **C**. Associated quantifications. We separately report the number of telomeric and γH2AX foci along with the colocalized events (Telomeric Induced Foci, TIFs) per nuclei in each isogenic clone (n=50 per quantification). Medians, quartiles and all data points are shown in violin plots. Holm-Sidak’s multiple comparison test; α = 0.05. *p* *** <0,001; *p* **** <0,0001.

**Supplemental Figure 2.**
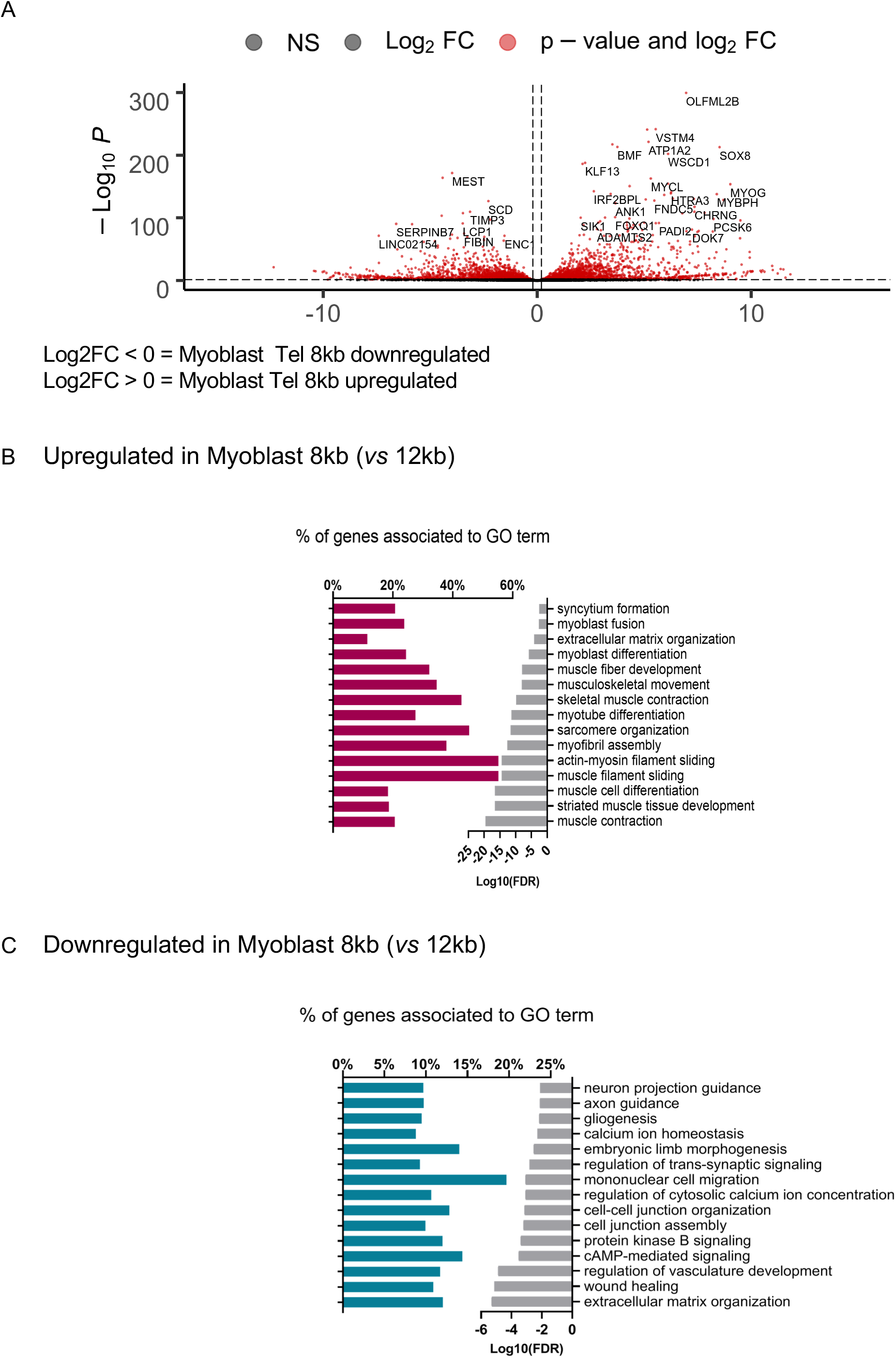
**A**. Volcano plots associated with transcriptomic analysis (RNASeq) for differential expression comparing isogenic clones of myoblasts with long (12kb) and shorter (8kb) telomeres. Log2(FC) and −log10 (FDR) are plotted on the *x-* and *y-*axis, respectively. Black dots represent genes that did not reach the significance thresholds whereas differentially expressed genes are shown in red. **B-C**. Gene Ontology (GO) for Biological pathways (BP) corresponding to enrichment analysis of upregulated (**B**) of downregulated (**C**) DEGs in myoblasts with short telomeres (8 kb) *vs*. myoblasts with long telomeres (12 kb) filtered on −2>FC>2 and FDR <0.05. Bar plots in the left (dark red/blue; respectively) represent the percentage DEG out of the total genes of associated with a GO-term shown in the right column. Grey bars in the right represent (Log10 of False Discovery Rate) for each BP.

**Supplemental Figure 3.**
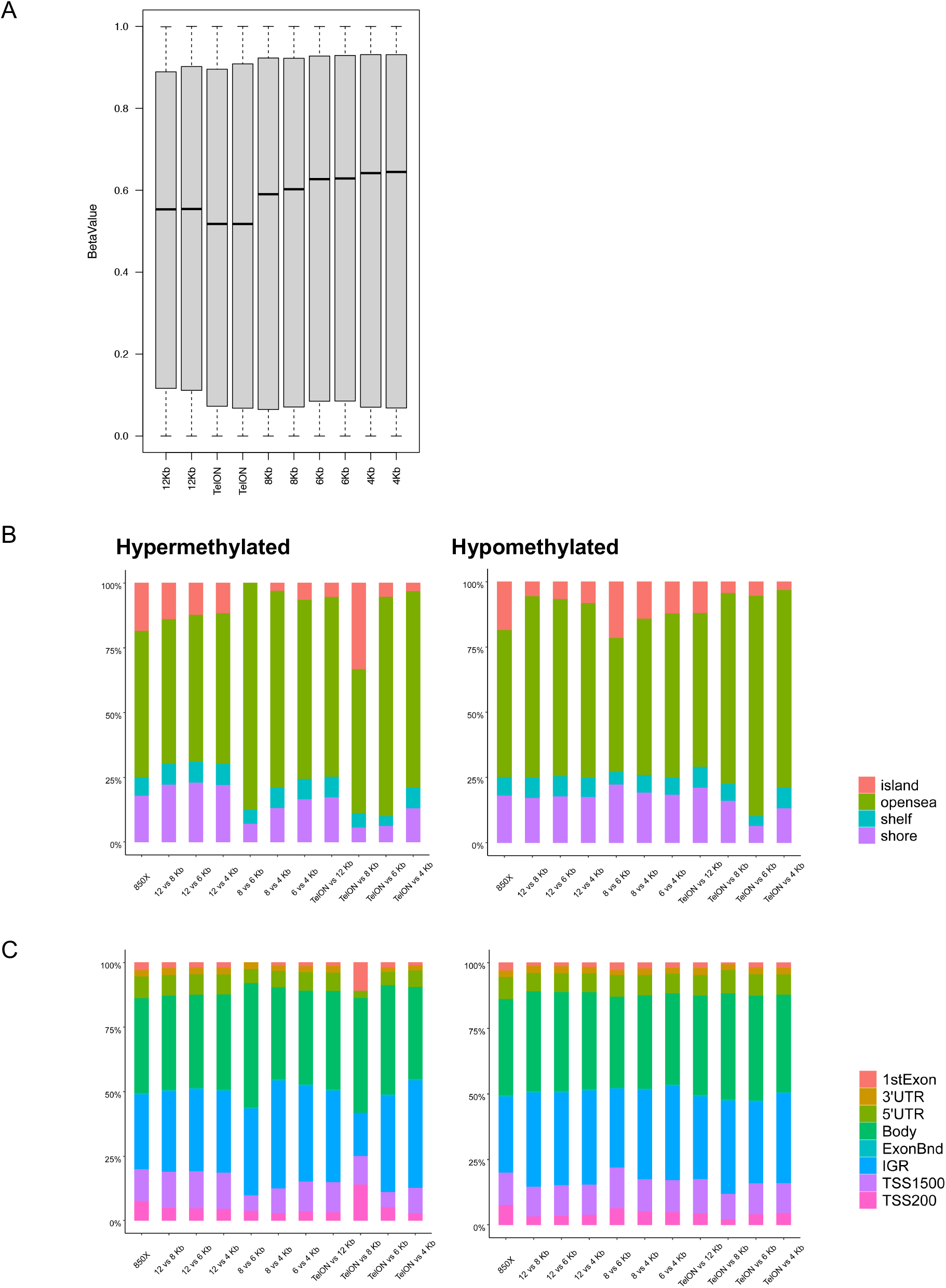
Boxplot reporting the median methylation mean from methylome array (EpicArray 850K) in myoblasts with different telomere length. TelON corresponds to myoblasts where the telomerase was not removed. **B**. Stacked barplots representing the distribution of Hypermethylated (left) and Hypomethylated (right) probes relative to CpG islands, shores (2kb flanking CpG islands), shelves (2kb extending from shores) or openseas (isolated CpG in the rest of the genome) in myoblast with shorter telomere (8kb) compared to isogenic clones with long telomeres (12kb; padj < 0.05 and abs (deltaBeta) > 0.2). **C**. Stacked barplots representing the distribution of hypermethylated (left) and hypomethylated (right) probes corresponding to genes first exon; 3’ UTR; 5’UTR; gene bodies; exon boundaries; internal genomic regions (IGR); probes located 1500 bp from transcription start sites (TSS1500) or 200 bp from transcription start sites (TSS200) in myoblast with shorter telomeres (8kb) compared to isogenic clones with long telomeres (12kb; padj < 0.05 and abs (deltaBeta) > 0.2).

**Supplemental Figure 4.**
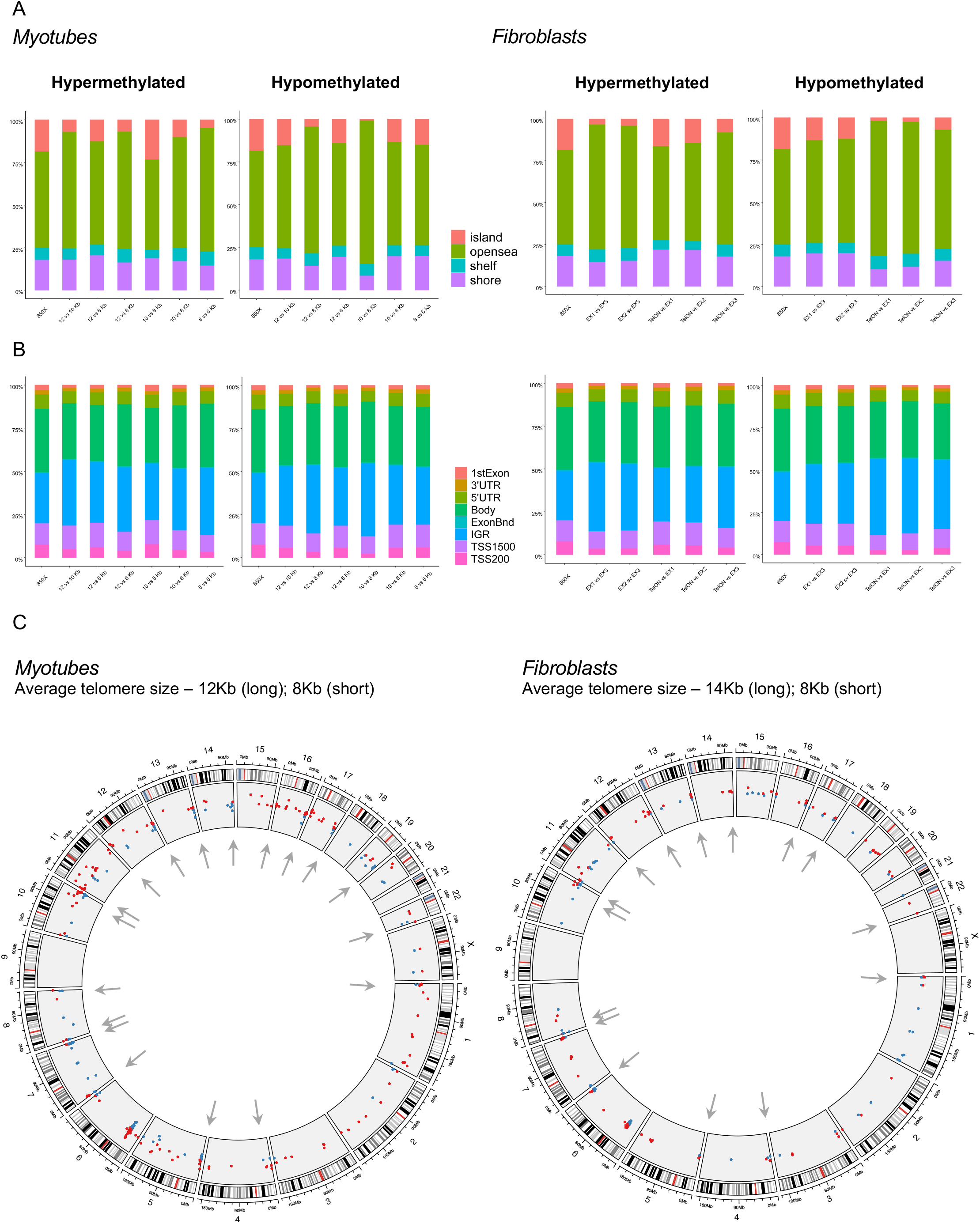
**A**. Stacked barplots representing the distribution of Hypermethylated and Hypomethylated probes relative to CpG islands, shores (2kb flanking CpG islands), shelves (2kb extending from shores) or openseas (isolated CpG in the rest of the genome) in either myotubes with shorter telomeres (8kb) compared to isogenic clones with long telomeres (left) or fibroblasts with shorter telomeres compared to isogenic clones with long telomeres (14kb, right). padj < 0.05 and abs (deltaBeta) > 0.2). **B**. Stacked barplots representing the distribution of hypermethylated and hypomethylated probes corresponding to genes first exon; 3’ UTR; 5’UTR; gene bodies; exon boundaries; internal genomic regions (IGR); probes located 1500 bp from transcription start sites (TSS1500) or 200 bp from transcription start sites (TSS200) in either myotubes with shorter telomeres (8kb) compared to isogenic clones with long telomeres (left) or fibroblasts with shorter telomeres (8kb) compared to isogenic clones with long telomeres (11kb, right). padj < 0.05 and abs (deltaBeta) > 0.2). **C**. Distribution of differentially methylated regions (DMRs) across the genome in either isogenic myotubes clones with long (12kb) and shorter telomere (8kb) or isogenic fibroblasts clones with long and shorter telomeres (left, right; respectively). Arrows (grey) point to DMRs located at subtelomeres.

**Supplemental Figure 5.**
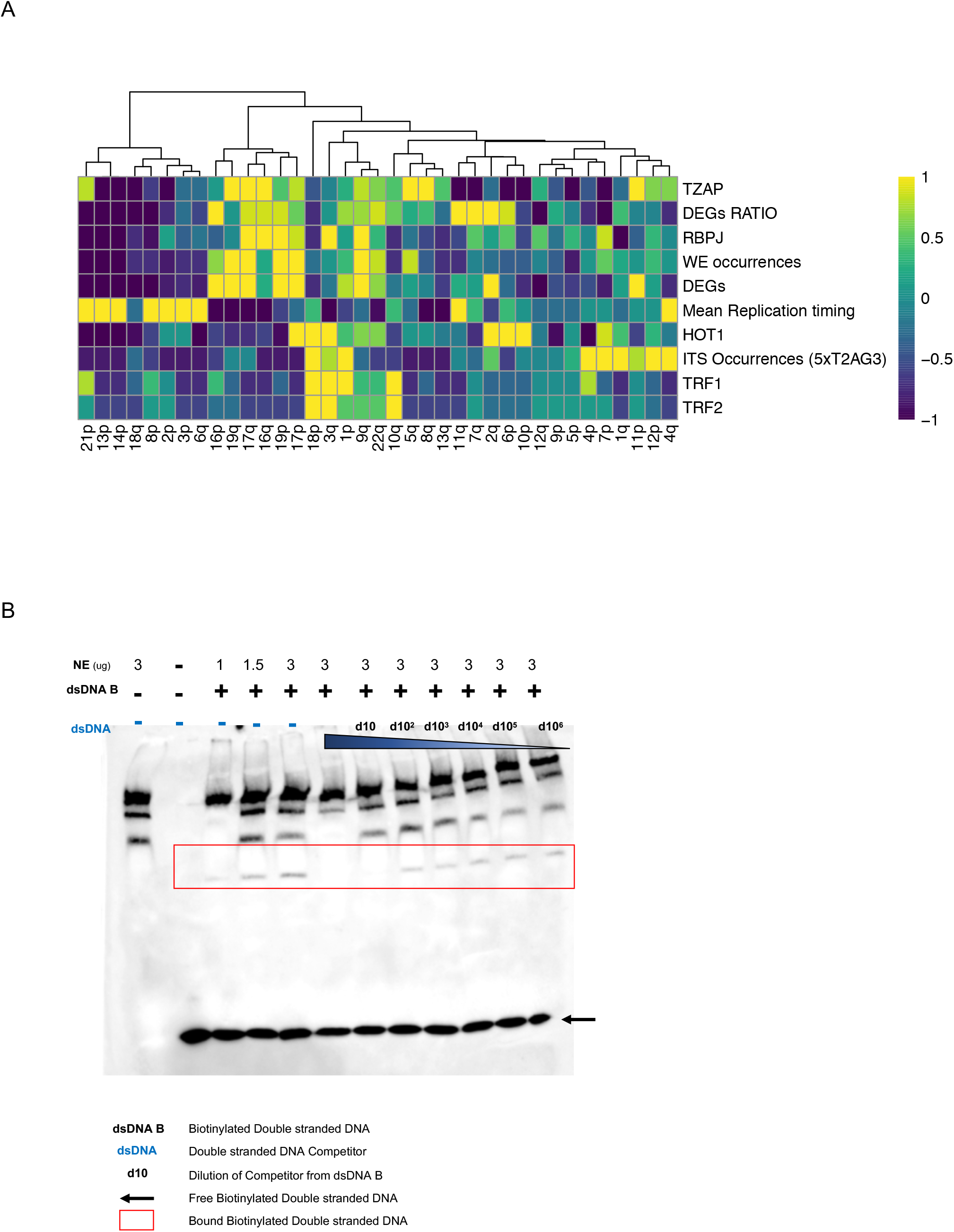
**A**. Unsupervised hierarchical clustering (WardD2; Manhattan distance) of subtelomeres (10Mb) with motif occurrences (WE), DEGs and telomeres associated factors (replication timing, proteins, Internal Telomeric Sequences ITS). Each parameter was extracted from available data set and normalized to a mean of 1. We report DEGs as total DEGs localized at each subtelomere and DEGs ratio when corrected for the gene density. **B**. Electrophoretic mobility shift assay (EMSA) results for WE 3’-CCTCCCAAAGTGCTGGGATTACAGGCGTGAGCCAC-5’ binding (dsDNA B). 5’ biotin labeled WE dsDNA was titrated with increasing amount of Nuclear Extract (NE) and unlabeled dsDNA competitor (blue) using serial dilutions. Biotinylated labeled dsDNA associated to a protein complex is shown in the red square; free unbound biotinylated dsDNA by a black arrow.

**Supplemental Figure 6.**
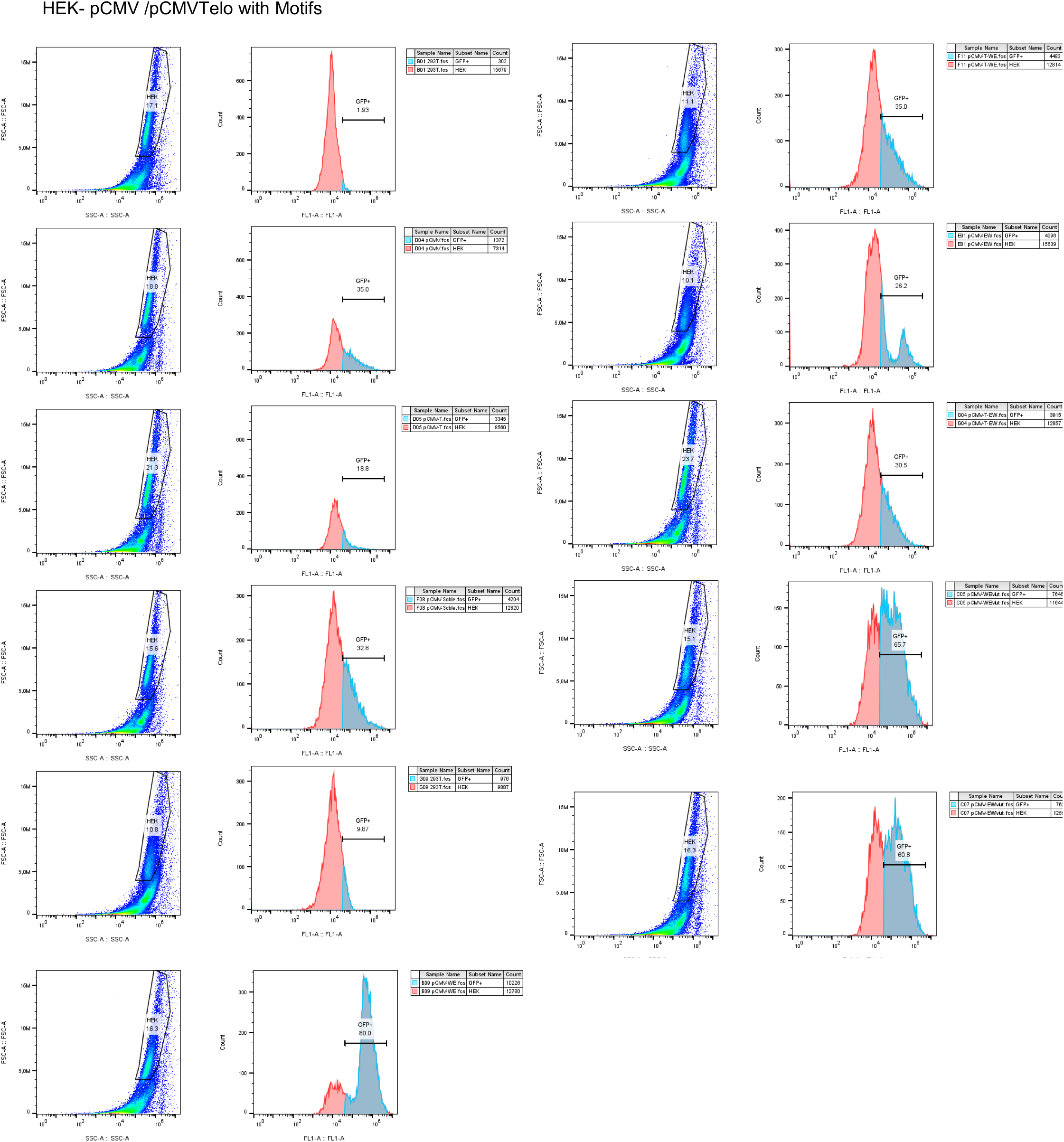
Representative flow cytometry plots of cells (HEK 293T) stably transfected with respective constructs of dsDNA motifs (WE, EW, Scble, WEMut, EWMut) cloned in a reporter vector without (pCMV) or with (pCMVTelo) a telomere seed used for testing position and telomeric position effect, respectively. We report the complete flow chart of live cells sorted using FSC-A and SSC-A parameters (left) along with histograms (right) reporting the proportion of eGFP positive cells (FL1-A; in blue) within the selected population. The threshold of eGFP detection was determined using untransfected HEK 293T(eGFP-negative cells). The same gates (population, eGFP+ cells) were kept for all analyses. Associated quantifications are reported in Figure 3.

**Supplemental Figure 7.**
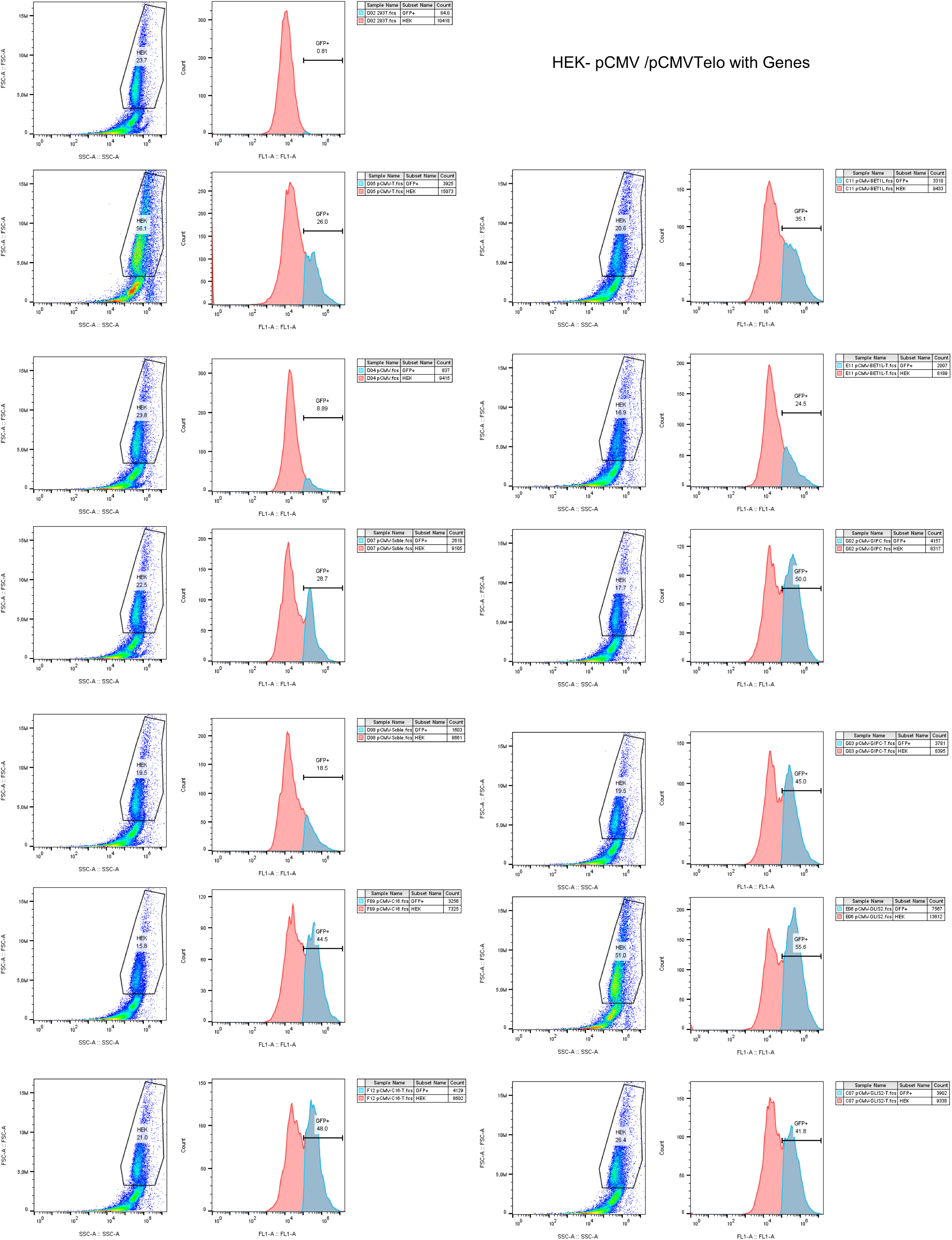
Representative flow cytometry plots of cells (HEK 293T) stably transfected with respective constructs containing dsDNA motifs found in direct proximity of DEGs from our transcriptomic analysis (*C16Orf*; *BET1L*; *GIPC3*; *GLIS2*) cloned in a reporter vector without (pCMV) or with (pCMVTelo) a telomere seed for testing position and telomeric position effect, respectively. We report the complete flow chart of cells sorted using FSC-A and SSC-A parameters (left) along with histograms (right) reporting the proportion of eGFP positive cells (FL1-A; in blue) with the complete selected population (live cells). Negative eGFP cells were determined by using untransfected HEK 293T cells. The same gates (population, eGFP+) were kept all analyses. Associated quantifications are reported in Figure 4.

**Supplemental Figure 8.**
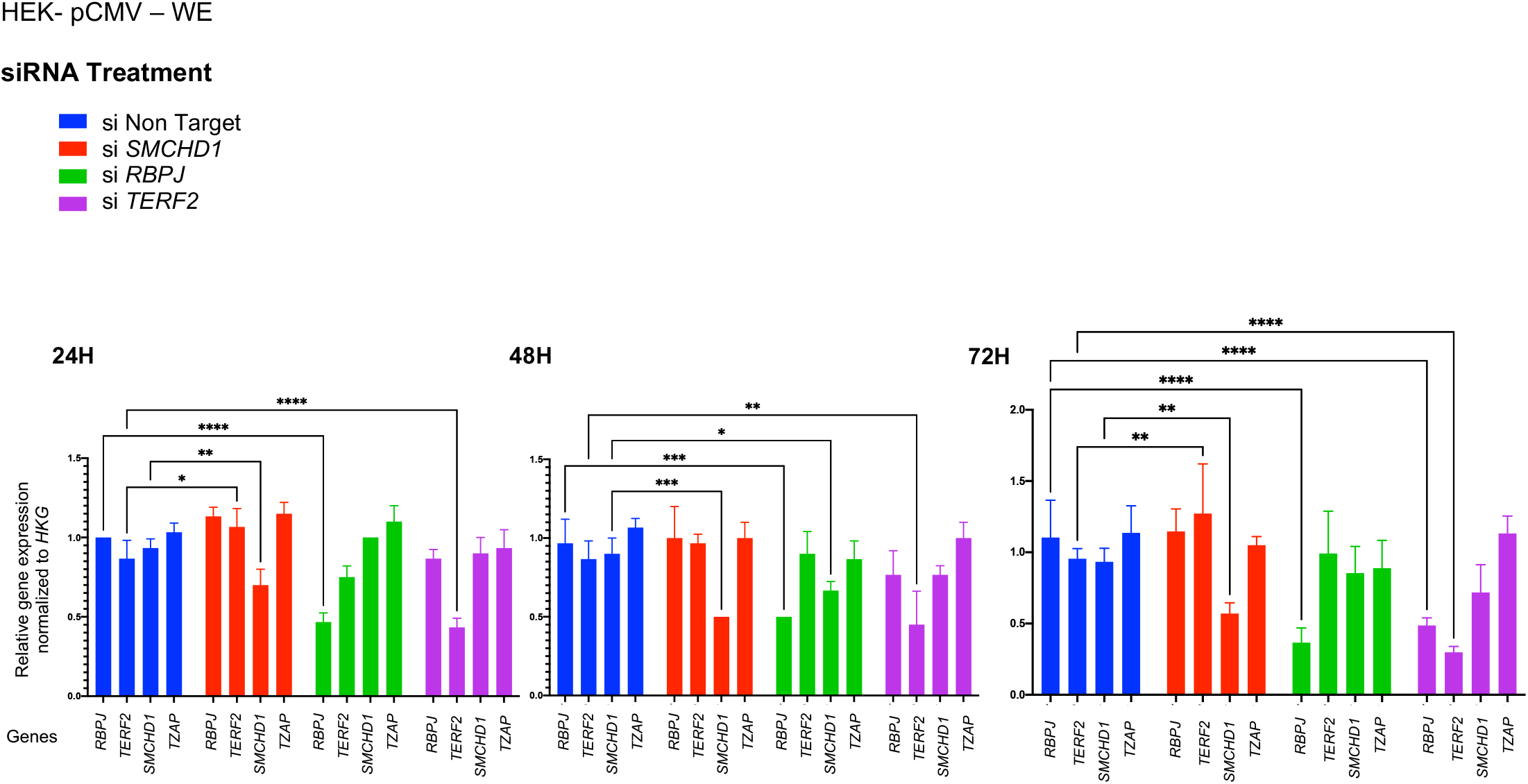
Relative gene expression (RT-qPCR) in HEK 293T cells transfected with siRNAs targeting *SMCHD1, RBPJ* and *TERF2*; respectively. Expression of selected genes (*RBPJ, TERF2, SMCHD1, TZAP*) is normalized to housekeeping genes (HKG; *PPIA, HPRT, GAPDH*) and respective expression at 24, 48 and 72 hours in the Non-targeted siRNA condition. For each condition, we report the average of three independent assays with technical duplicates. Mean ± SEM are shown. Holm-Sidak’s multiple comparison test; α = 0.05. *p* * <0,05; *p* ** <0,005; *p* *** <0,001; *p* **** <0,0001.

**Supplemental Figure 9.**
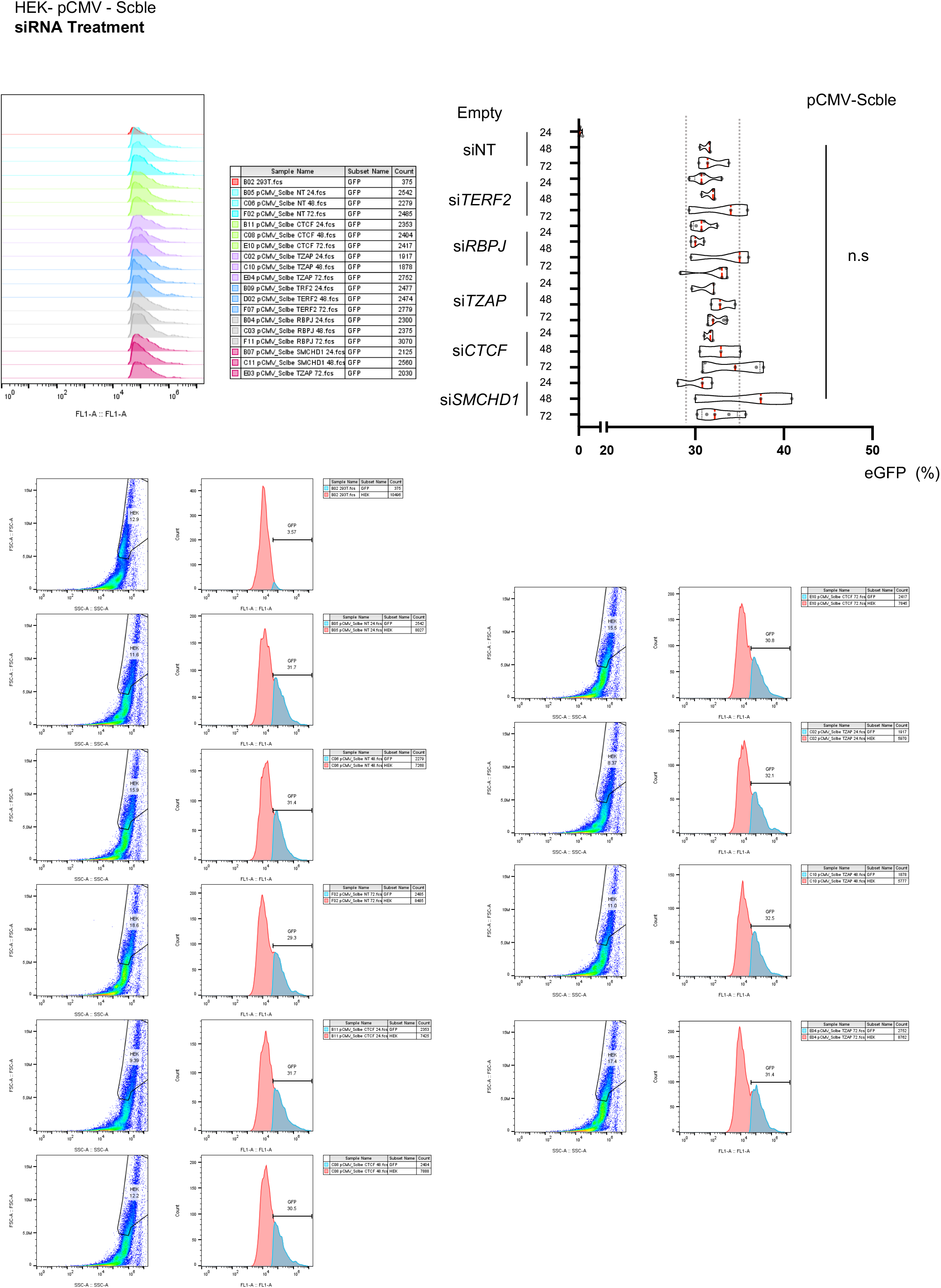
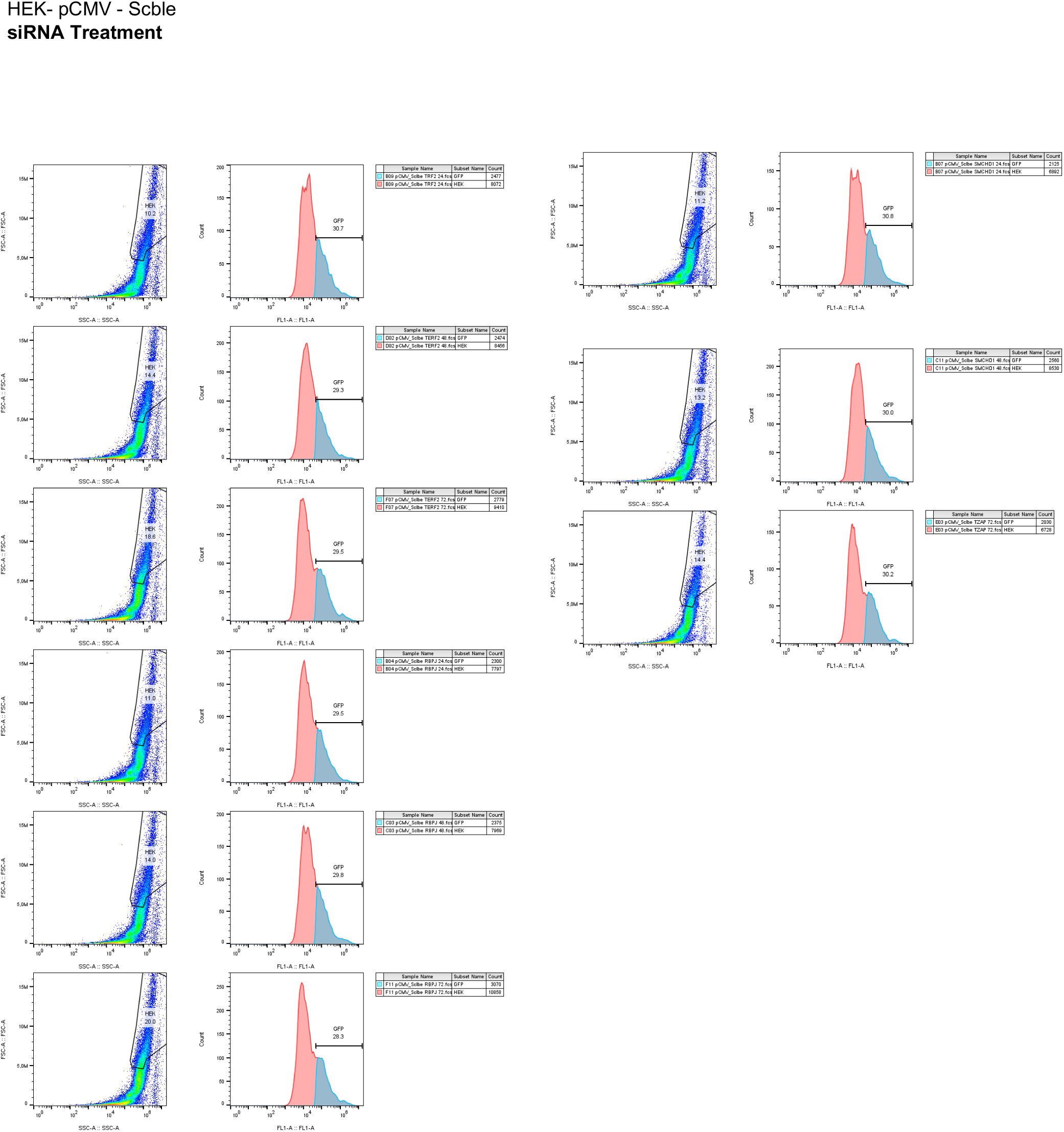
Representative flow cytometry plots of cells (HEK 293T) stably transfected with a dsDNA motif (Scble condition, as described in Figure 2.) inserted in a pCMV reporter plasmid and transfected with siRNAs along with their associated quantifications. We report the complete flow charts of cells sorted using FSC-A and SSC-A parameters (left) along with histograms (right) reporting the proportion of eGFP-positive cells (FL1-A; in blue) at 24, 48 and 72 hours post treatments. Negative eGFP cells were determined by using untransfected HEK 293T cells. The same gates (population, eGFP+) were kept for all analyses. For each condition, we report biological quadruplicate (n=4) in box and violins. Medians and quartiles are shown (red and dashed lines, respectively); dashed grey lines are shown to represent the variability associated with the NT condition. Holm-Sidak’s multiple comparison test; α = 0.05.

**Supplemental Figure 10.**
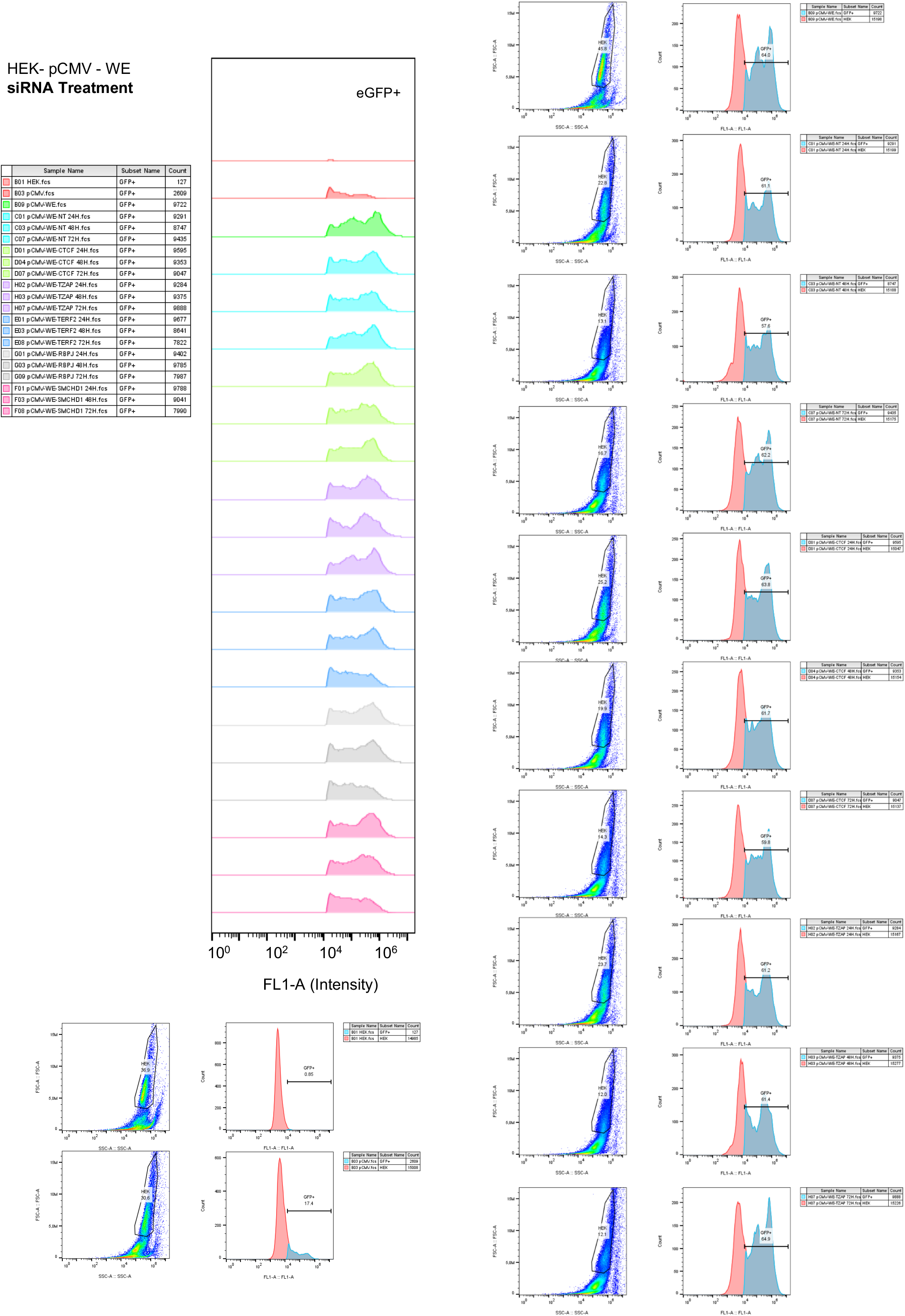
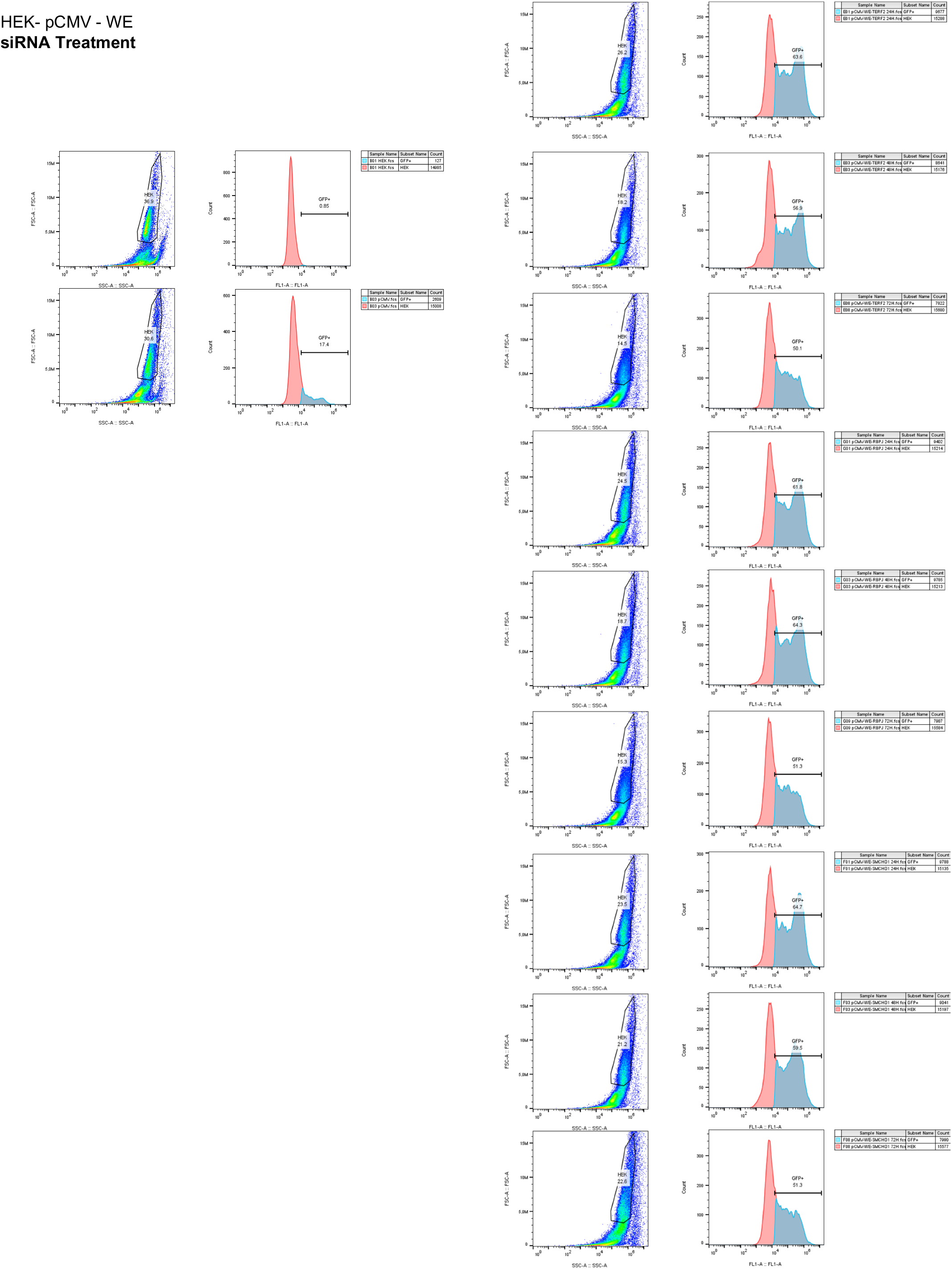
Representative flow cytometry plots of cells (HEK 293T) stably transfected with a dsDNA motif (WE) inserted in a pCMV reporter plasmid and treated with siRNAs. We report the complete flow charts of cells sorted using FSC-A and SSC-A parameters (left) along with histograms (right) reporting the proportion of eGFP positive cells (FL1-A; in blue) for the selected population at 24, 48 and 72 hours post treatments. Negative eGFP cells were determined by using untransfected HEK 293T cells without transfection. The same gates (population, eGFP+) were kept for all of analyses. Associated quantifications are reported in Figure 4.

**Supplemental Figure 11.**
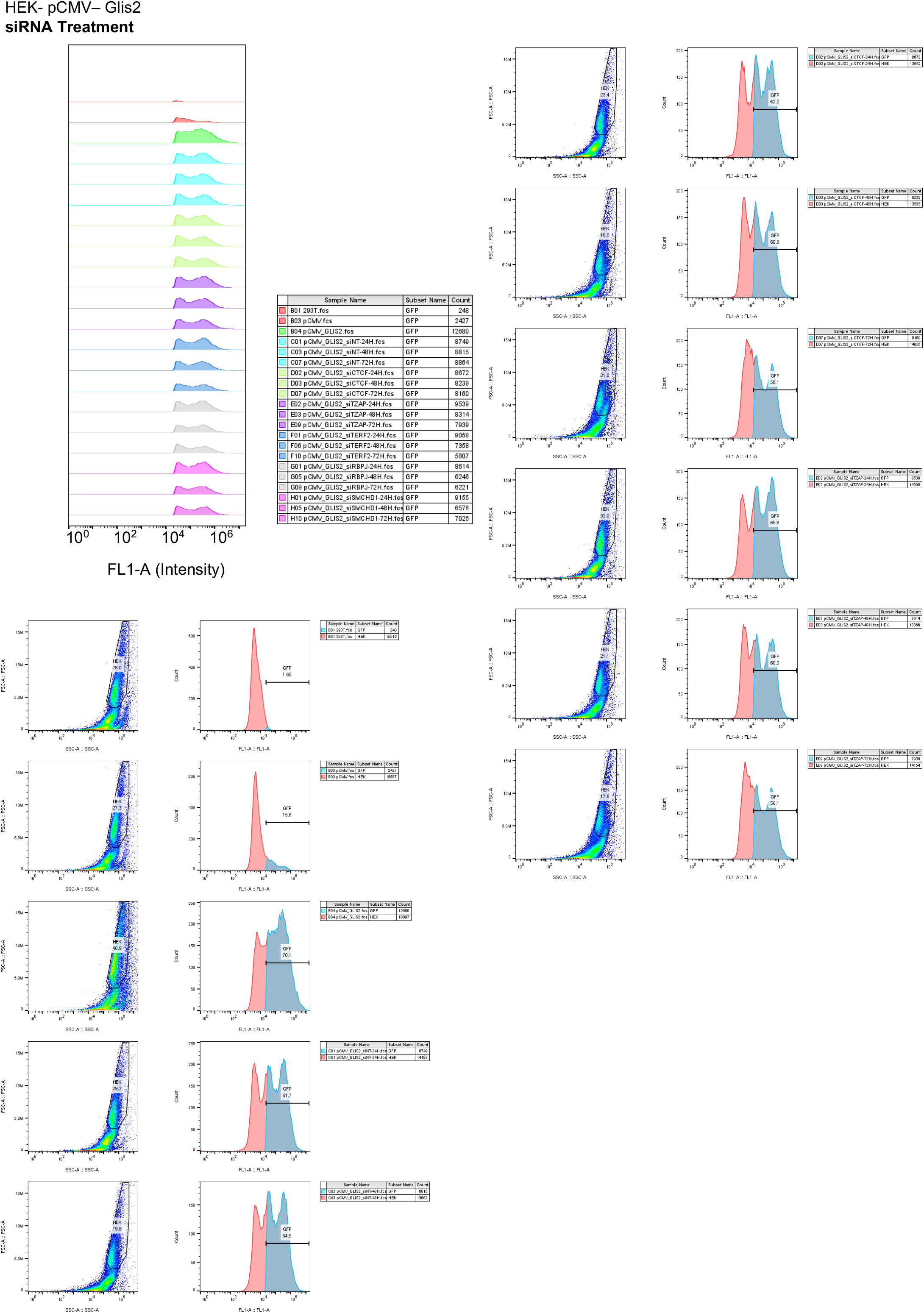
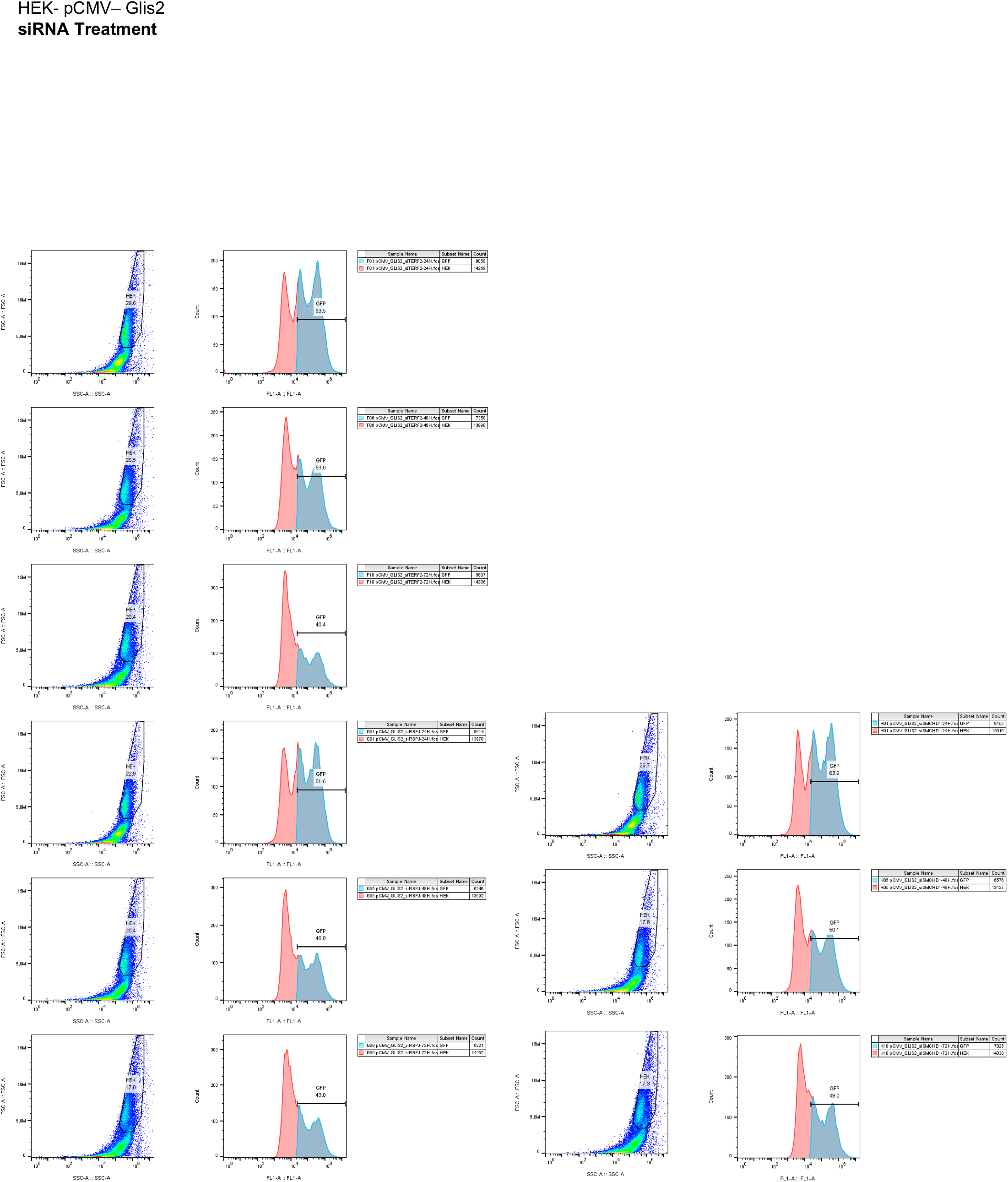
Representative flow cytometry plots of cells (HEK 293T) stably transfected with a dsDNA motif (*Glis2*) inserted in a pCMV reporter plasmid and treated with siRNAs. We report the complete flow charts of cells sorted using FSC-A and SSC-A parameters (left) along with histograms (right) reporting the proportion of eGFP positive cells (FL1-A; in blue) for the selected population at 24, 48 and 72 hours post treatments. Negative eGFP cells were determined by using untransfected HEK 293T. The same gates (population, eGFP+) were kept for all analyses. Associated quantifications are reported in Figure 4.

**Supplemental Figure 12.**
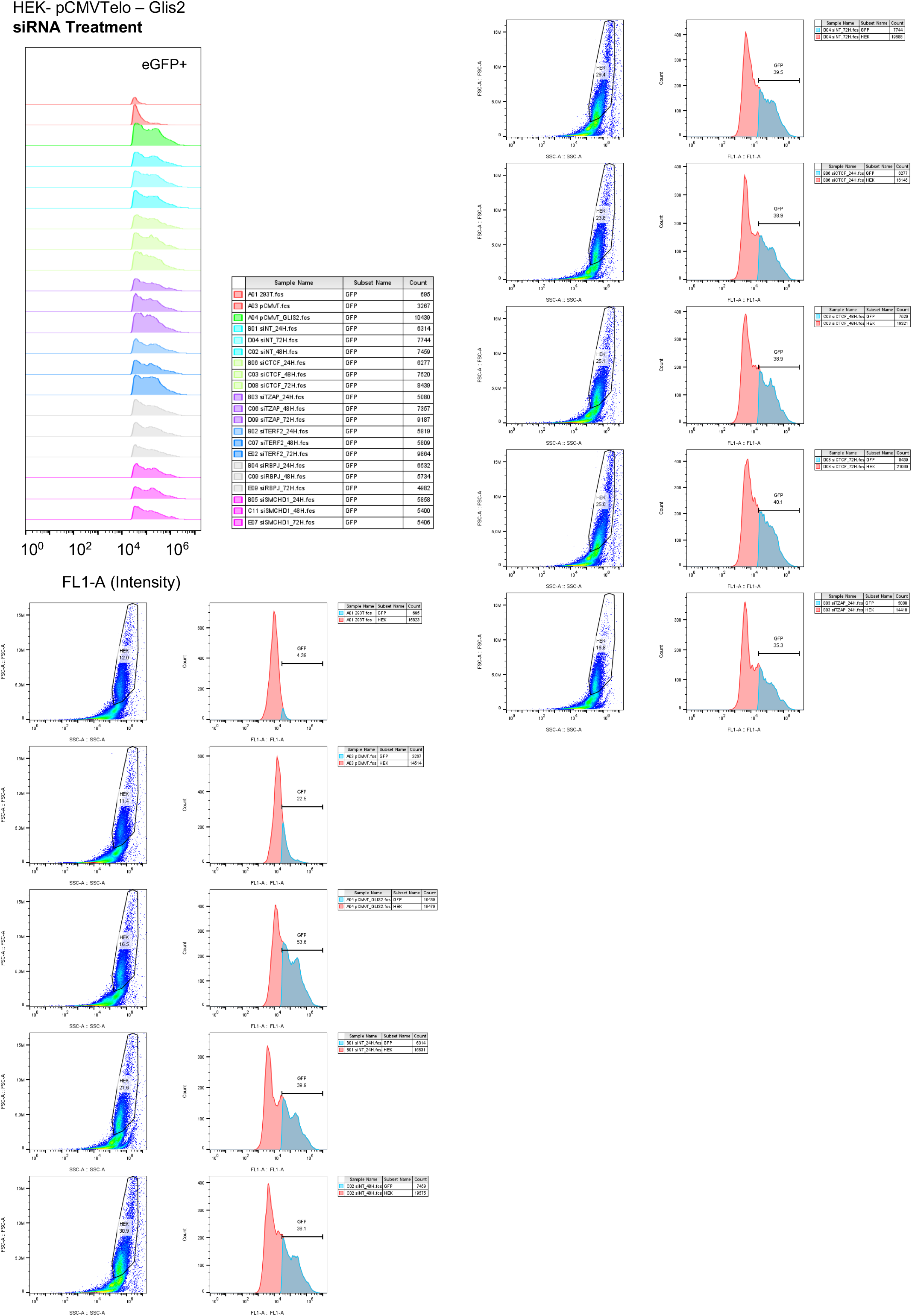
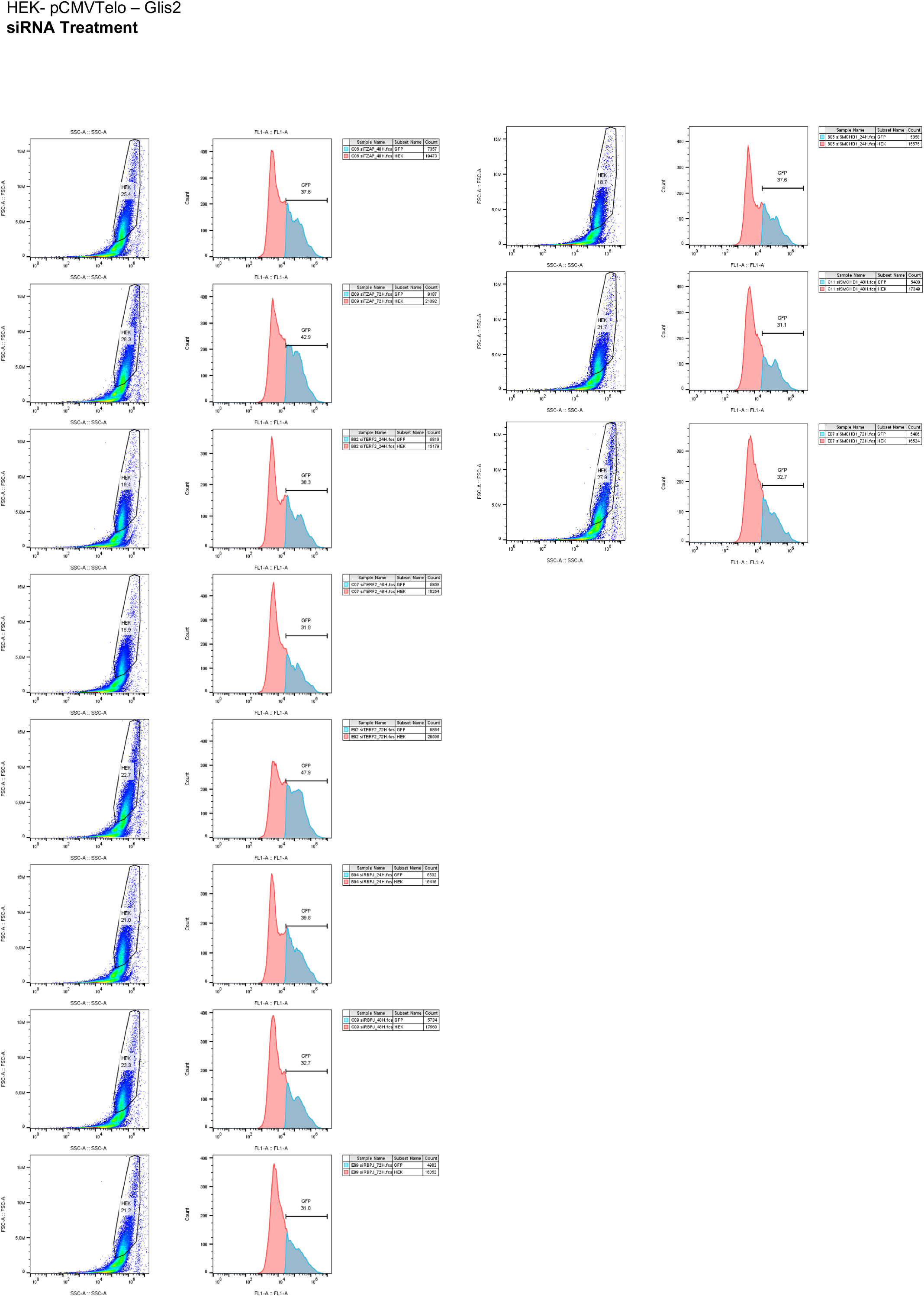
Representative flow cytometry plots of cells (HEK 293T) stably transfected with a dsDNA motif (Glis2) inserted in a pCMVTelo (with a telomere seed) reporter plasmid and treated with siRNAs. We report the complete flow charts of cells sorted using FSC-A and SSC-A parameters (left) along with histograms (right) reporting the proportion of eGFP positive cells (FL1-A; in blue) for the selected population at 24, 48 and 72 hours post treatments. Negative eGFP cells were determined by using untransfected HEK 293T. The same gates (population, eGFP+) were kept all analyses. Associated quantifications are reported in Figure 4.

**Supplemental Figure 13.**
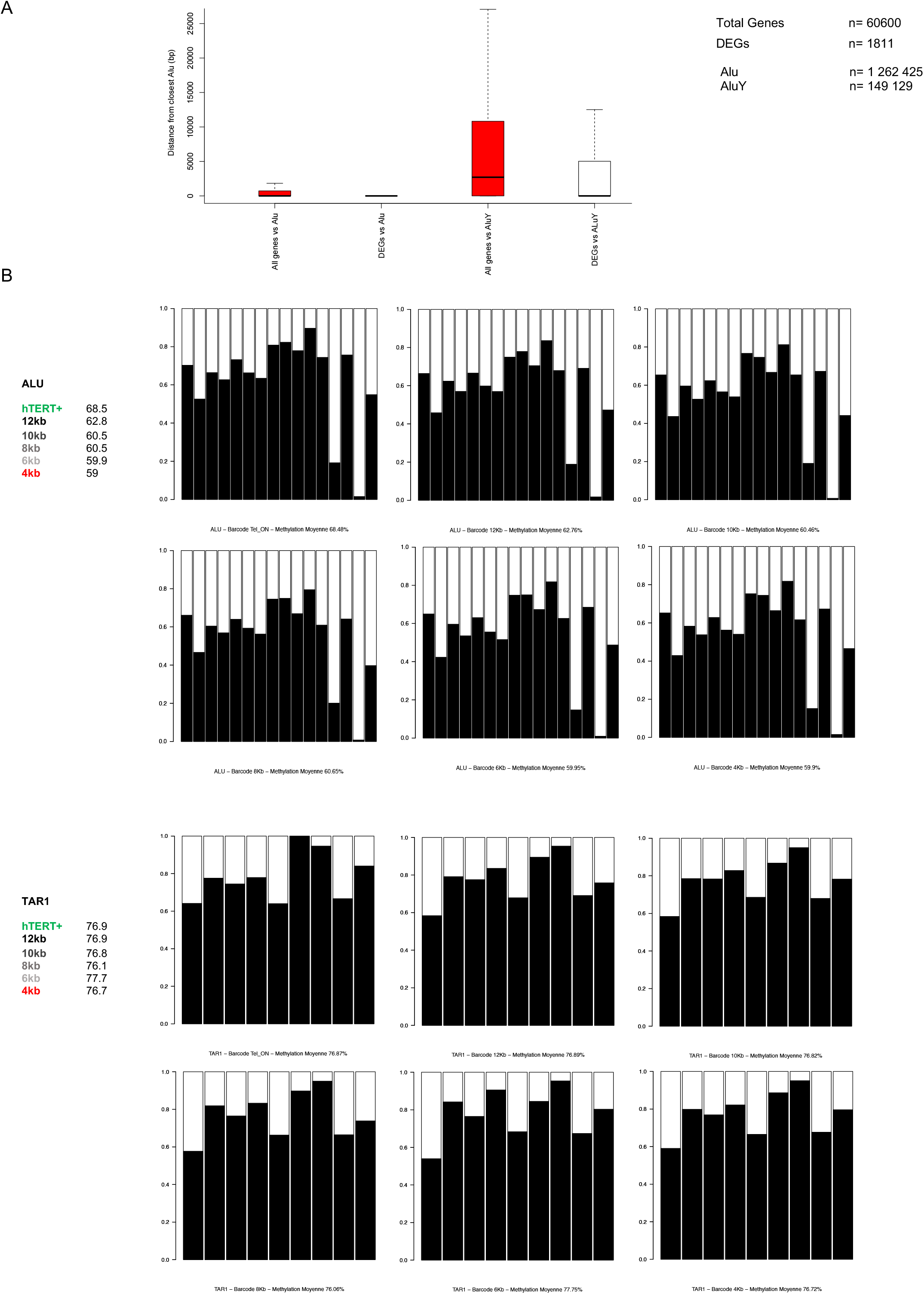
**A**. Distance of genes and DEGs to the closest Alu or AluY elements. Distances were calculated using the total number of genes (n=60 600) or DEGs from our experiments (n=1811) and reference location of either Alu (n= 1 262 425) or AluY (n= 149 129). For Alu comparison; Welch Two Sample t-test, *p-value* < 2.2e-16; for AluY, Welch Two Sample t-test, p-value < 2.2e-16. **B**. Stacked barplots representing the distribution of methylation at CpGs in either Alu repeats (top panel) or TAR1 repeats (bottom panel) in isogenic clones of myoblasts with long and shorter telomeres. Methylation was assessed using sodium bisulfited-converted DNA and Bisulfite sequencing primers. A minimum of 50 000 sequences were aligned per condition.

**Supplemental Figure 14.**
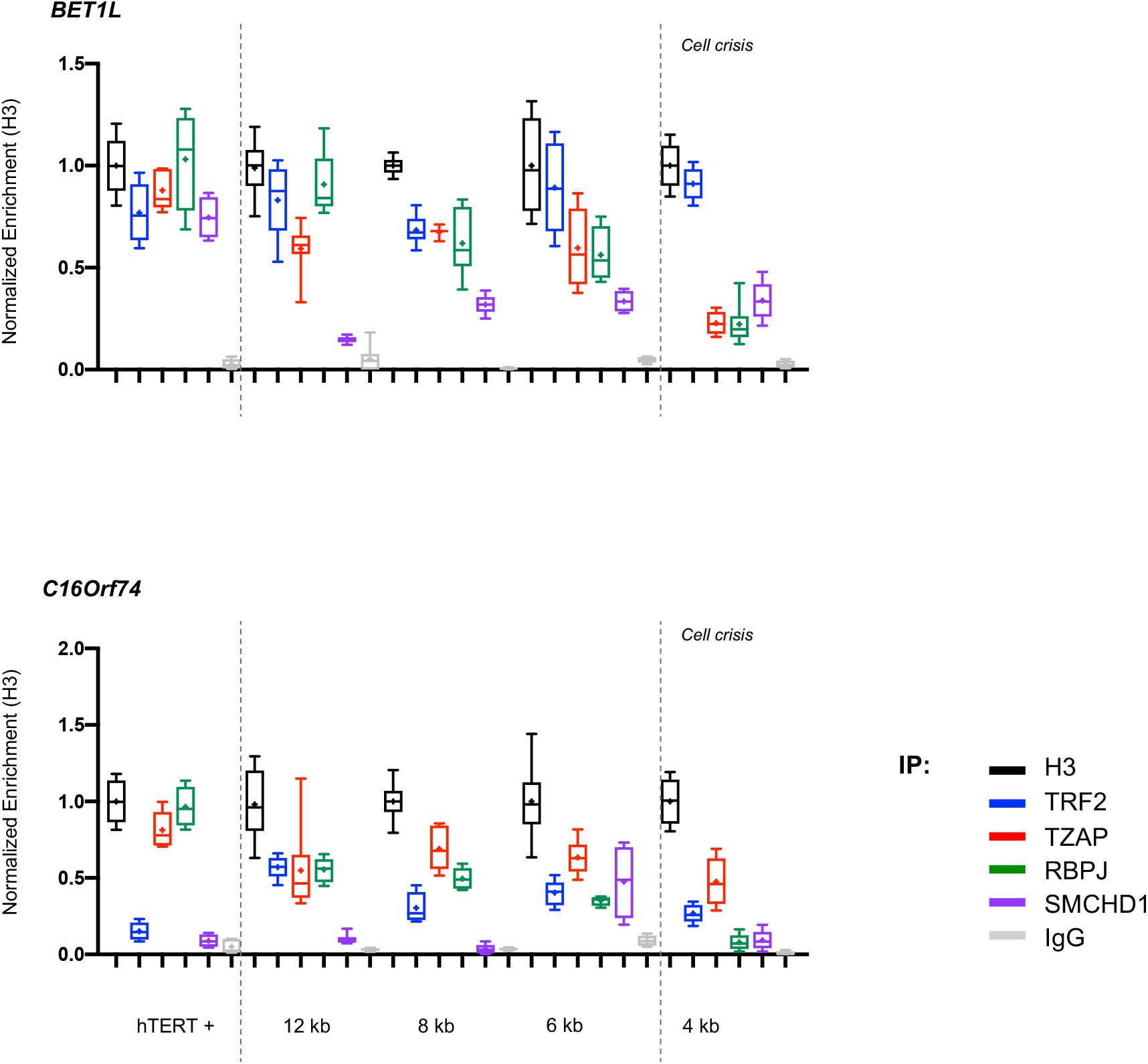
Protein enrichment at *BET1L* and *C16Orf74* loci detected by ddPCR after ChIP in isogenic clones with various telomere lengths. We report the enrichment of H3 (black), TRF2 (blue), TZAP (red), RBPJ (green), SMCHD1(purple) and IgG (grey) normalized to H3. Dashed grey lines are shown to represent the critical telomere length, before cell crisis (active telomerase, 4kb; respectively). ChIP-ddPCR were performed in biological quadruplicate. Holm-Sidak’s multiple comparison test; α = 0.05. *p* * <0,05; *p* ** <0,005; *p* *** <0,001; *p* **** <0,0001.

## Notes

### Competing Interest Statement

The authors have declared no competing interest.

## References

1. de Lange, T. Shelterin: the protein complex that shapes and safeguards human telomeres. Genes Dev. 19, 2100–2110 (2005).

2. Azzalin, C. M., Reichenbach, P., Khoriauli, L., Giulotto, E. & Lingner, J. Telomeric repeat containing RNA and RNA surveillance factors at mammalian chromosome ends. Science 318, 798–801 (2007).

3. Feretzaki, M., Renck Nunes, P. & Lingner, J. Expression and differential regulation of human TERRA at several chromosome ends. RNA 25, 1470–1480 (2019).

4. Lim, C. J. et al. The structure of human CST reveals a decameric assembly bound to telomeric DNA. Science 368, 1081–1085 (2020).

5. Denchi, E. L. & de Lange, T. Protection of telomeres through independent control of ATM and ATR by TRF2 and POT1. Nature 448, 1068–1071 (2007).

6. Galati, A. et al. TRF1 and TRF2 binding to telomeres is modulated by nucleosomal organization. Nucleic Acids Res. 43, 5824–5837 (2015).

7. Takai, H., Smogorzewska, A. & de Lange, T. DNA Damage Foci at Dysfunctional Telomeres. Current Biology 13, 1549–1556 (2003).

8. Demanelis, K. et al. Determinants of telomere length across human tissues. Science 369, eaaz6876 (2020).

9. Daniali, L. et al. Telomeres shorten at equivalent rates in somatic tissues of adults. Nat Commun 4, 1597 (2013).

10. Bendix, L., Horn, P. B., Jensen, U. B., Rubelj, I. & Kolvraa, S. The load of short telomeres, estimated by a new method, Universal STELA, correlates with number of senescent cells. Aging Cell 9, 383–397 (2010).

11. Zou, Y., Sfeir, A., Gryaznov, S. M., Shay, J. W. & Wright, W. E. Does a sentinel or a subset of short telomeres determine replicative senescence? Mol Biol Cell 15, 3709–3718 (2004).

12. López-Otín, C., Blasco, M. A., Partridge, L., Serrano, M. & Kroemer, G. The hallmarks of aging. Cell 153, 1194–1217 (2013).

13. Shay, J. W. & Wright, W. E. Role of telomeres and telomerase in cancer. Semin. Cancer Biol. 21, 349–353 (2011).

14. Counter, C. M. et al. Telomere shortening associated with chromosome instability is arrested in immortal cells which express telomerase activity. EMBO J. 11, 1921–1929 (1992).

15. Grolimund, L. et al. A quantitative telomeric chromatin isolation protocol identifies different telomeric states. Nat Commun 4, 2848 (2013).

16. Kan, S. L., Saksouk, N. & Déjardin, J. Proteome Characterization of a Chromatin Locus Using the Proteomics of Isolated Chromatin Segments Approach. Methods Mol. Biol. 1550, 19–33 (2017).

17. Déjardin, J. & Kingston, R. E. Purification of Proteins Associated with Specific Genomic Loci. Cell 136, 175–186 (2009).

18. Bottoni, G. et al. CSL controls telomere maintenance and genome stability in human dermal fibroblasts. Nat Commun 10, 3884 (2019).

19. Kappei, D. et al. HOT1 is a mammalian direct telomere repeat-binding protein contributing to telomerase recruitment. EMBO J. 32, 1681–1701 (2013).

20. Gauchier, M., van Mierlo, G., Vermeulen, M. & Déjardin, J. Purification and enrichment of specific chromatin loci. Nat. Methods 14, 986–10 (2020).

21. Jahn, A. et al. ZBTB48 is both a vertebrate telomere-binding protein and a transcriptional activator. EMBO Rep. 18, 929–946 (2017).

22. Li, J. S. Z. et al. TZAP: A telomere-associated protein involved in telomere length control. Science 355, 638–641 (2017).

23. Johnson, J. E. & Macdonald, R. J. Notch-independent functions of CSL. Curr Top Dev Biol 97, 55–74 (2011).

24. Maicas, M., Jimeno-Martín, Á., Millán-Trejo, A., Alkema, M. J. & Flames, N. The transcription factor LAG-1/CSL plays a Notch-independent role in controlling terminal differentiation, fate maintenance, and plasticity of serotonergic chemosensory neurons. PLoS Biol. 19, e3001334 (2021).

25. Laberthonnière, C., Magdinier, F. & Robin, J. D. Bring It to an End: Does Telomeres Size Matter? Cells 8, 30 (2019).

26. de Bruin, D., Kantrow, S. M., Liberatore, R. A. & Zakian, V. A. Telomere folding is required for the stable maintenance of telomere position effects in yeast. Mol. Cell. Biol. 20, 7991–8000 (2000).

27. Baur, J. A., Zou, Y., Shay, J. W. & Wright, W. E. Telomere position effect in human cells. Science 292, 2075–2077 (2001).

28. Stadler, G. et al. Telomere position effect regulates DUX4 in human facioscapulohumeral muscular dystrophy. Nature Structural & Molecular Biology 20, 671–678 (2013).

29. Robin, J. D. et al. Telomere position effect: regulation of gene expression with progressive telomere shortening over long distances. Genes Dev. 28, 2464–2476 (2014).

30. Robin, J. D. & Magdinier, F. Physiological and Pathological Aging Affects Chromatin Dynamics, Structure and Function at the Nuclear Edge. Front Genet 7, 153 (2016).

31. Dong, X. et al. Age-related telomere attrition causes aberrant gene expression in sub-telomeric regions. Aging Cell 20, e13357 (2021).

32. Mukherjee, A. K. et al. Telomere length-dependent transcription and epigenetic modifications in promoters remote from telomere ends. PLOS Genetics 14, e1007782 (2018).

33. Robin, J. D. et al. SORBS2 transcription is activated by telomere position effect-over long distance upon telomere shortening in muscle cells from patients with facioscapulohumeral dystrophy. Genome Res. 25, 1781–1790 (2015).

34. Bork, S. et al. DNA methylation pattern changes upon long-term culture and aging of human mesenchymal stromal cells. Aging Cell 9, 54–63 (2010).

35. Franzen, J. et al. DNA methylation changes during long-term in vitro cell culture are caused by epigenetic drift. Commun Biol 4, 598 (2021).

36. Bigot, A. et al. Age-Associated Methylation Suppresses SPRY1, Leading to a Failure of Re-quiescence and Loss of the Reserve Stem Cell Pool in Elderly Muscle. Cell Reports 13, 1172–1182 (2015).

37. Kim, W. et al. Regulation of the Human Telomerase Gene TERT by Telomere Position Effect-Over Long Distances (TPE-OLD): Implications for Aging and Cancer. PLoS Biol. 14, e2000016 (2016).

38. Robin, J. D. et al. Mitochondrial function in skeletal myofibers is controlled by a TRF2-SIRT3 axis over lifetime. Aging Cell 19, e13097 (2020).

39. Simonet, T. et al. The human TTAGGG repeat factors 1 and 2 bind to a subset of interstitial telomeric sequences and satellite repeats. Cell Res. 21, 1028–1038 (2011).

40. Hammal, F., de Langen, P., Bergon, A., Lopez, F. & Ballester, B. ReMap 2022: a database of Human, Mouse, Drosophila and Arabidopsis regulatory regions from an integrative analysis of DNA-binding sequencing experiments. Nucleic Acids Res. 50, D316–D325 (2022).

41. Arnoult, N. et al. Replication timing of human telomeres is chromosome arm-specific, influenced by subtelomeric structures and connected to nuclear localization. PLOS Genetics 6, e1000920 (2010).

42. Piqueret-Stephan, L., Ricoul, M., Hempel, W. M. & Sabatier, L. Replication Timing of Human Telomeres is Conserved during Immortalization and Influenced by Respective Subtelomeres. Sci Rep 6, 32510–13 (2016).

43. Koering, C. E. et al. Human telomeric position effect is determined by chromosomal context and telomeric chromatin integrity. EMBO Rep. 3, 1055–1061 (2002).

44. Ottaviani, A. et al. Identification of a perinuclear positioning element in human subtelomeres that requires A-type lamins and CTCF. EMBO J. 28, 2428–2436 (2009).

45. Vančevska, A. et al. SMCHD1 promotes ATM-dependent DNA damage signaling and repair of uncapped telomeres. EMBO J. 39, e102668 (2020).

46. Young, E., Abid, H. Z., Kwok, P.-Y., Riethman, H. & Xiao, M. Comprehensive Analysis of Human Subtelomeres by Whole Genome Mapping. PLOS Genetics 16, e1008347 (2020).

47. Horvath, S. DNA methylation age of human tissues and cell types. 14, R115–20 (2013).

48. Bell, C. G. et al. DNA methylation aging clocks: challenges and recommendations. 20, 249–24 (2019).

49. Wildschutte, J. H., Baron, A., Diroff, N. M. & Kidd, J. M. Discovery and characterization of Alu repeat sequences via precise local read assembly. Nucleic Acids Res. 43, 10292–10307 (2015).

50. Grover, D., Mukerji, M., Bhatnagar, P., Kannan, K. & Brahmachari, S. K. Alu repeat analysis in the complete human genome: trends and variations with respect to genomic composition. Bioinformatics 20, 813–817 (2004).

51. Dridi, S. Alu mobile elements: from junk DNA to genomic gems. Scientifica (Cairo) 2012, 545328 (2012).

52. Ferrari, R. et al. TFIIIC Binding to Alu Elements Controls Gene Expression via Chromatin Looping and Histone Acetylation. Molecular Cell 77, 475–487.e11 (2020).

53. Su, M., Han, D., Boyd-Kirkup, J., Yu, X. & Han, J.-D. J. Evolution of Alu elements toward enhancers. Cell Reports 7, 376–385 (2014).

54. Payer, L. M. et al. Alu insertion variants alter gene transcript levels. Genome Res. 31, 2236–2248 (2021).

55. Jintaridth, P. & Mutirangura, A. Distinctive patterns of age-dependent hypomethylation in interspersed repetitive sequences. Physiol Genomics 41, 194–200 (2010).

56. Bollati, V. et al. DNA methylation in repetitive elements and Alzheimer disease. Brain Behav Immun 25, 1078–1083 (2011).

57. Dion, C. et al. SMCHD1 is involved in de novo methylation of the DUX4-encoding D4Z4 macrosatellite. Nucleic Acids Res. 40, 663 (2019).

58. Laberthonnière, C. et al. SMCHD1 variants may induce variegated expression in Facio Scapulo Humeral Dystophy and Bosma Arhinia and microphtalmia syndrome. bioRxiv 2021.05.17.444338 (2021).

59. Castel, D. et al. Dynamic binding of RBPJ is determined by Notch signaling status. Genes Dev. 27, 1059–1071 (2013).

60. Wang, H. et al. NOTCH1-RBPJ complexes drive target gene expression through dynamic interactions with superenhancers. Proc. Natl. Acad. Sci. U.S.A. 111, 705–710 (2014).

61. Kappei, D. & Londoño-Vallejo, J. A. Telomere length inheritance and aging. Mechanisms of Ageing and Development 129, 17–26 (2008).

