## Supplementary material for "Telomere Position Effect Over Long Distance acts as a genome-wide epigenetic regulator through a common *cis*- element": Material-Methods

### Materials and Methods

#### Cell Culture

Myoblasts and fibroblasts were immortalized with a floxable hTERT cassette as described previously[1,2]. The human embryonic kidney 293T cell line (CRL-1573) was obtained from ATCC. For day-to-day maintenance, human myoblasts were seeded in dishes coated with 0.1% pigskin gelatin in 4:1 Dulbecco modified Eagle medium/Medium 199 supplemented with 15% FBS, 0.02M HEPES, 1.4mg/l vitamin B12, 0.03mg/l ZnSO4, 0.055mg/l dexamethasone, 2.5µg/l hepatocyte growth factor and 10µg/l basic fibroblast growth factor. Myogenicity was verified by myotubes formation following a change to differentiation medium (2% horse serum in 4:1 Dulbecco modified Eagle medium: Medium 199) when 70-90% confluent.

Fibroblasts and HEK293 cells were grown in DMEM with L-alanyl-L-glutamine (GlutaMAX TM I), D-glucose and sodium pyruvate (Life technologies). Media were supplemented in with 10% fetal bovine serum (FBS, Gibco).

All cultures were maintained in a 5% oxygen environment and passaged at ~60% confluency. Population doublings (PDs) were calculated as PD = ln[(final number of cells)/(initial number of cells)]/ln(2).

#### Telomere Restriction Fragment analysis (TRF)

Terminal restriction fragment assay was done as previously described with a slight modification, using a biotin-labelled probe[2].

#### Telomeric Induced Foci (TIFs).

Cells were grown on cover slides and fixed for 10min on ice with 4% paraformaldehyde in 1x PBS. After PBS washes (3x5min), cells were incubated for 1h at room temperature (RT) in blocking solution (1%Triton X-100, 1%BSA, 5% donkey serum in PBS). To perform the PNA-FISH staining, cells were washed twice with SSC2X for 5min at RT and subsequently treated with RNase A for 45min at 37°C. After an additional SSC2X wash (5min, 4°C), coverslides were dried and incubated upside-down with a hybridization solution containing the PNA probe (20µl H2O; 70µl formamide; 7µl 10% blocking B (Roche); 1µl 1M Tris pH7.2; 1µl probe) and sealed on coverslips using rubber cement. Slides were then heated at 85°C for 4min and incubated in the dark at 37°C in a humidification chamber for 2 hours. After removal of the rubber cement, cells were serially washed in three different solutions: twice for 15min at RT with washing solution I (10mM Tris pH 7.2; 70% formamide); twice for 15min at RT with washing solution II (150mM NaCl; 50mM Tris pH7.2; 0.05% Tween 20) and twice with 1x PBS for 5min at RT. Cells were then blocked for 1 hour with the blocking solution and immuno-stained overnight at 4°C in the blocking solution containing the primary rabbit polyclonal anti-53BP1 antibody (1:500; Novus Biologicals). After three washes with PBS/0.1% Triton X-100, slides were incubated for 1h30min at RT with Alexa 555 Donkey anti-rabbit secondary antibody in PBS containing 0.5% Triton X-100, 1% BSA, 2.5% donkey serum. Slides were mounted in Vectashield with DAPI (Vector Laboratories, Burlingame, USA). Images were taken using a Confocal system (LSM800, Zeiss). Co-localization events, representing telomeric DNA damages (TIFs), were counted in at least 30 nuclei per condition from three independent experiments using the IMARIS software.

#### DNA extraction and Sodium bisulfite sequencing

DNA was extracted from the different types of samples using the NucleoSpin Tissue (Macherey-Nagel) according to manufacturer's instructions. For sodium bisulfite sequencing,

1µg of genomic DNA was denatured for 30 minutes at 37°C in NaOH 0.4N and incubated overnight in a solution of 3M Sodium bisulfite pH5 and 10mM Hydroquinone using previously described protocol[3]. Converted DNA was purified using the Wizard DNA CleanUp kit (Promega) following manufacturer's recommendation and precipitated by ethanol precipitation for 5 hours at -20°C. Primers were designed in order to amplify methylated and unmethylated DNA with the same efficiency using the MethPrimer software[4], avoiding the presence of CpGs in the primer sequence. After deep sequencing, (>50K sequences retrieved), sense and antisense sequences were assembled in a single sequence and bam file converted to fastq file. After trimming of each BSP primers, data were aligned using the BiQ Analyser HiMod software (http://biq-analyzer.bioinf.mpi-inf.mpg.de)[5] and processed in R (version 3.4.2). BiQ Analyzer HiMod converts sequencing data by using « 1 » for a methylated CG, « 0 » for unmethylated and « x » in case of misalignment. For each sequenced fragment, three methylation score are calculated, (i) the CpG methylation score of each CpG, (ii) the sequence methylation score, corresponding to the average methylation level of each sequence and (iii) the global methylation score that corresponds to the global level of methylation for each biological sample in a given region calculated as the ratio of methylated CpG with the number of aligned CpG for all sequences and CpG for a given biological sample as previously described[6].

**Infinium MethylationEPIC Array**

Genome-wide DNA methylation analysis was performed by using the Infinium MethylationEPIC Array through Diagenode services. Genomic DNA was extracted using the NucleoSpin Tissue kit (Macherey-Nagel) from two different cell pellets for each sample. A minimum of 500ng was sent to Diagenode services for DNA methylation analysis. The analysis was mainly carried out using the ChAMP R package[7]. Probes with missing values are removed, samples with more than 10% of probes with a detection p-value of greater than 0.01, probes with a detection p-value of greater than 0.01 in at least one sample and probes for which 5% of samples have a bead count less than 3 are filtered out. An annotation file is loaded that contains information about the location of probes - such as chromosome, position and nearby genes [8]. Probes targeting CpG sites that are near SNPs, or align with multiple locations are filtered out. A matrix of Methylation Beta values is returned. The Beta-value indicates the percentage of copies of a CpG site from a given sample that were methylated[9] determined for each CpG location as the relative intensity of methylated signal (M) and unmethylated signal (U)[10]. Samples from the different groups were compared to identify Differentially Methylated Probes (DMPs) based on a significantly different average methylation level at CpG site. The ChAMP function for identifying DMPs uses the limma package[11]. An adjusted p-value < 0.05, and absolute value of difference in mean Beta value > 0.2, were required for a probe to be identified as a DMP [12]. A OLS linear model is fit to the methylation values of each probe to estimate the effect of belonging to a phenotype group vs the control group, Moderated t-statistics are used to calculate p-values, the p-values are adjusted with the B&H method. The Probe Lasso method was used to identify regions of differential methylation (Differentially Methylated Regions, DMR) using the champ DMR function. ProbeLasso starts by selecting DMPs (default p-vlaue cutoff = 0.05) and extending a lasso in either direction. The size of the lasso depends on the gene feature (Body, IGR etc) and CGI relation (Island, Shore, etc) associated with probe location. The rationale for differing lasso sizes is that probe density varies between gene features and CGI relations. Mini DMRs are created when the number of DMPs contained within a lasso is greater or equal to a minimum value. Overlapping or neighboring mini DMRs are merged until the distance between merged DMRs is at least 1,000 bp. A p-value is calculated for each DMR, using Stouffer's weighted method to combine the p-values of the probes contained in the DMR. The default cut off p-value for DMRs to be selected is 0.05, and they must contain at least 7 probes and exceed 50 bp in width. The Combat method was used to correct for batch effects[13] . Heatmaps were realized using the pheatmap (v1.0.12) R package using Beta value matrix for methylation. DMPs with an FDR adjusted pvalue < 0.05 and an abs(deltaBeta) > 0.2 were represented as barplot using the ggplot2 (v3.3.3) R package for genomic features and CGI status according to Illumina Epic array annotation file. Genomic localizations of DMRs were represented as circos using Circulize v0.4.15. NCBI

Gene Expression Omnibus (https://www-ncbi-nlm-nih-gov.gate2.inist.fr/geo/) DNA methylation data accession number: XXXXXX accessible using the XXXXXX

**RNA Sequencing.**

***RNA extraction, quality control and library preparation***

Total RNA was extracted using the RNAeasy kit (Qiagen) following manufacturer's instructions. Quality, quantification and sizing of total RNA was evaluated using the RNA 6000 Pico assay (Agilent Technologies Ref. 5067-1513) on an Agilent 2100 Bioanalyzer system. The RNA integrity number (RIN) was calculated for each sample and only samples with a RIN >9 were kept for further use. Libraries were constructed using 2 µg of total RNA. The TruSeq Stranded mRNA Library Preparation Kit High Throughput (Illumina, ref RS-122-2103) was used according to the manufacturer's guidelines. Briefly, PolyA+ containing RNA molecules were purified using polyT oligo-attached magnetic beads. Thermal fragmentation was carried out after two rounds of enrichment for PolyA+ mRNA. cDNA was synthesized using reverse transcriptase (Superscript IV) and random primers. This was followed by second strand cDNA synthesis, end repair process, adenylation of 3' ends and ligation of the adapters. The products were then purified and enriched with 15 cycles of PCR to create the cDNA library. Libraries were quantified by qPCR using the KAPA Library Quantification Kit for Illumnia Libraries (Roche, ref. 7960140001). Library profiles were assessed using the DNA High Sensitivity LabChip Kit (Agilent Technologies Ref. 5067-4626) on an Agilent Bioanalyzer 2100. Libraries were sequenced on an Illumina Next-seq 500 2x75bp at the GBiM genomic core facilities (https://www.marseille-medical-genetics.org/fr/genomics-bioinformatics-platform/).

***RNA-Seq data processing and differential expression analysis***

We assessed fastq sequence data quality using FastQC v0.11.5 and trimmed the reads to remove adapter sequences and low-quality bases using Trimmomatic v0.36. The resulting trimmed single-end reads were aligned using STAR v2.5.3a to the GRCh38 human genome release. Obtained BAM files were indexed using Sambamba (v0.6.6) after ordering them by coordinates.

Aligned reads were counted with StringTie v1.3.1c using GENCODE annotation. DEGs of different conditions were identified using R package DESeq2 (v1.18.1) with the following thresholds FDR < 0,05, abs (LogFC) >2.

***DEGs analysis***

DEGs were represented as circos using Circlize v0.4.15 for visualization.

Overrepresentation test analyses were performed using enrichGO from the R package clusterProfiler (v3.10.15). With DEGs as input, we identified biological processes (BP) with an FDR< 0.05 using a custom gene universe based on genes with a rowmean for counts > 1. Results are presented as barplot with on the bottom x-axis, the log10(FDR), corresponding GO terms and on the top x-axis, the percentage of DEGs associated with a GO-term.

##### ***Heatmap***

Heatmaps were realized using R package pheatmap (v1.0.12) with unsupervised hierarchical clustering scaled on rows. Input used either TPM counts for RNA-seq or custom ratios for comparison of multi sourced data.

#### ***De novo*** **motif analysis**

Motif analysis was performed using MEME (v5.1.1) on identified DMRs, CHIP-seq peaks from available datasets (TRF1 and TRF2, GSE26005; TZAP, GSE96778; HOT1, GSE46237; RBPJ, GSE29498) or at DEGs coordinates using fasta file. Motif reported were selected for an Evalue < 0.05 using markov order 1 (for bias in CG) parameters for DMRs.

*meme file.fa -o output -dna -mod anr -nmotifs 10 -evt 0.05 -minw 6 -maxw 35 -markov_order 0/1 -V -maxsize 20000000 -p 10*

#### Nuclear protein extraction

Nuclear protein extraction was performed using NE-PER Nuclear and cytoplasmic extraction (reagents #78835; Thermo Scientific)

#### Electrophoretic mobility shift assay

EMSA were performed using either 2 agarose or 10% polyacrylamide gels. Revelation for EMSA in polyacrylamide gel with biotin incorporated dsDNA fragment (WE) was performed using the LightShift™ Chemiluminescent EMSA Kit (ref 20148).

#### Transfection and Flow cytometry

pCMV derived plasmids are described in details in[14]. DNA fragments were cloned downstream of the eGFP reporter gene in pCMV vectors. DNA insert were obtained either from strand synthesis and hybridization for WE motif or PCR amplification for motif in genomic context and inserted at the *Asc*I restriction site. Details are available upon request. Sanger sequencing was performed for all selected clones containing the insert in a 5' to 3' orientation. Prior to transfection, linearization of vectors was achieved using the *BstX*I restriction enzyme (NEB). Transfection of the linearized vectors was performed using a modified calcium phosphate method[15] optimized in order to obtain a single integration per cell[15]. Three days post-transfection, the Hygromycin B selection antibiotics was added to the culture medium (Life technologies) at a final concentration of 400µg/ml. Cells were kept under permanent selection for several passages (3 weeks on average). At different time points, eGFP expression was analyzed using an Accuri flow cytometer and processed using the FlowJo software (Becton-Dickinson). The percentage of eGFP-positive cells was determined using the corresponding non-transfected cells as the baseline for autofluorescence. Mean values (M1) were used to compare fluorescence in the different samples.

#### 3D DNA FISH

Three-dimensional DNA Fluorescent in situ Hybridization was performed as previously described previously[2]. For HEK293 we used a probe generated by Nick translation using the pCMV or pCMV plasmid as template.

**siRNA transfections**

Transfections were performed using DharmaFECT1 Transfection reagents and siRNAs (SMARTpool ON-TARGETplus Human, 5nmol; Horizon discovery) targeting *TERF2* (ID: 7014); *RBPJ* (ID: 3516); *TZAP* (ID: 3104); *CTCF* (ID: 10664); *SMCHD1* (ID: 23347) and Control Non-Targeting, NT (D-001810-0X). Transfections were carried on following the protocol and conditions provided for HEK293 by the manufacturer (Horizon discovery).

**RT-qPCR.**

RNA extraction was realized using the Qiagen's RNeasy mini kit. Briefly, reverse transcription of 1µg of total RNA was performed using the Superscript IV kit and oligo dT following manufacturer's instructions (Life Technologies). Primers were designed using Primer Blast. PCR amplification was performed on a StepOnePlus Real-Time PCR system (Life Technologies) using the SYBR green master mix with the following program: Pre-incubation at 95°C for 10 minutes then 40 cycles amplification each corresponding to 15 seconds at 95°C followed by 1 minute at 60°C. The program ends with a melting step including a step of a second at 98°C, 30 seconds at 70°C and finally 10 seconds at 98°C.
Crossing-threshold (Ct) values were normalized by subtracting the geometric mean of three housekeeping genes (*GAPDH*, *PPIA* and *HPRT*). All Ct values were corrected by their PCR efficiency, determined by 1:2 or 1:4 cDNA dilution series. All analyzes were carried out in biological duplicates and technical duplicates. Primer sequences are provided as supplemental information.

**Chromatin Immunoprecipitation quantified by ddPCR (ChIP-ddPCR)**

Samples for chromatin immunoprecipitation (IP) were prepared as followed. IP either using H3 (Abcam ab1791), TZAP (GeneTex GTX 118671), RBPJ (Invitrogene 720219), SMCHD1 (equal mix Abcam ab31865 and Sigma HPA03944), TRF2 antibody (Imgenex124A) and IgG (Milipore PP64B) were crosslinked for 10 min at RT and 20 min at 4°C with 0.8% formaldehyde (methanol free, ultrapure EM grade, Polysciences, Inc; Warrington PA). Reaction was stopped at RT for 10 min with the addition of Glycine to a final concentration of 0.125 M. Cells were rinsed twice with ice-cold 1X PBS, scraped from the dish and pelleted by centrifugation (800g, 5min at 4°C). Next, cells were treated according to the manufacturer's instructions (Pierce Classic Protein G IP Kit, Thermo Scientific). For sonication, we used a total processing time of 15min per sample in a Bioruptor (Diagenode) using the following settings: 15 cycles; 30 Sec ON/30 Sec OFF on High power. Sonicated DNA was controlled on a 2% agarose gel; adequate sonication is achieved when a smear ranging from 200-700bp is obtained. IPs were processed using a 4°C O/N incubation (concentration of antibodies between 1.5 and 5µg according to manufacturer's instruction); 1µl of each preparation: IP, IgG, 1% input were used as controls for ddPCR analysis. Primers were designed for the WE-associated region of each gene; results are adjusted to inputs and further normalized to H3 IP. Each PCR primer pairs were tested on genomic DNA to verify specificity and efficiency.

**Statistical Analysis**

All experiments were repeated at least three times, with at least 3 biological replicates.

Quantitative data are displayed as means ± standard error of the mean or ± standard deviation when notified. Sample sizes as well as the statistical test used of each experiment are described in each corresponding figure legends or methods. Results from each group were treated with the GraphPad prism software for all statistical tests. All tests were two-sided and alpha set at 0.05. Only p-values less than 0.05 were considered statistically significant.

### References


1. Stadler, G. *et al.* Telomere position effect regulates DUX4 in human facioscapulohumeral muscular dystrophy. *Nature Structural & Molecular Biology* **20,** 671–678 (2013).
2. Robin, J. D. *et al.* Telomere position effect: regulation of gene expression with progressive telomere shortening over long distances. *Genes Dev.* **28,** 2464–2476 (2014).
3. Magdinier, F. *et al.* Regional methylation of the 5' end CpG island of BRCA1 is associated with reduced gene expression in human somatic cells. *FASEB J* **14,** 1585–1594 (2000).
4. Li, L.-C. & Dahiya, R. MethPrimer: designing primers for methylation PCRs. *Bioinformatics* **18,** 1427–1431 (2002).
5. Bock, C. *et al.* BiQ Analyzer: visualization and quality control for DNA methylation data from bisulfite sequencing. *Bioinformatics* **21,** 4067–4068 (2005).
6. Roche, S. *et al.* Methylation hotspots evidenced by deep sequencing in patients with facioscapulohumeral dystrophy and mosaicism. *Neurol Genet* **5,** e372 (2019).
7. Morris, T. J. *et al.* ChAMP: 450k Chip Analysis Methylation Pipeline. *Bioinformatics* **30,** 428–430 (2014).
8. Butcher, L. M. & Beck, S. Probe Lasso: a novel method to rope in differentially methylated regions with 450K DNA methylation data. *Methods* **72,** 21–28 (2015).
9. Du, P. *et al.* Comparison of Beta-value and M-value methods for quantifying methylation levels by microarray analysis. *BMC Bioinformatics* **11,** 587–9 (2010).
10. Bibikova, M. *et al.* High density DNA methylation array with single CpG site resolution. *Genomics* **98,** 288–295 (2011).
11. Ritchie, M. E. *et al.* limma powers differential expression analyses for RNA-sequencing and microarray studies. *Nucleic Acids Res.* **43,** e47 (2015).
12. Smyth, G. K. Linear models and empirical bayes methods for assessing differential expression in microarray experiments. *Stat Appl Genet Mol Biol* **3,** Article3 (2004).
13. Johnson, W. E., Li, C. & Rabinovic, A. Adjusting batch effects in microarray expression data using empirical Bayes methods. *Biostatistics* **8,** 118–127 (2007).
14. Ottaviani, A. *et al.* Identification of a perinuclear positioning element in human subtelomeres that requires A-type lamins and CTCF. *EMBO J.* **28,** 2428–2436 (2009).
15. Koering, C. E. *et al.* Human telomeric position effect is determined by chromosomal context and telomeric chromatin integrity. *EMBO Rep.* **3,** 1055–

1061 (2002).
